# Metagenomics of the modern and historical human oral microbiome with phylogenetic studies on *Streptococcus mutans* and *Streptococcus sobrinus*

**DOI:** 10.1101/842542

**Authors:** Mark Achtman, Zhemin Zhou

## Abstract

We have recently developed bioinformatic tools to accurately assign metagenomic sequence reads to microbial taxa: SPARSE [1] for probabilistic, taxonomic classification of sequence reads, EToKi [2] for assembling and polishing genomes from short read sequences, and GrapeTree [3], a graphic visualizer of genetic distances between large numbers of genomes. Together, these methods support comparative analyses of genomes from ancient skeletons and modern humans [2,4]. Here we illustrate these capabilities with 784 samples from historical dental calculus, modern saliva and modern dental plaque. The analyses revealed 1591 microbial species within the oral microbiome. We anticipated that the oral complexes of Socransky *et al.* [5] would predominate among taxa whose frequencies differed by source. However, although some species discriminated between sources, we could not confirm the existence of the complexes. The results also illustrate further functionality of our pipelines with two species that are associated with dental caries, *Streptococcus mutans* and *Streptococcus sobrinus.* They were rare in historical dental calculus but common in modern plaque, and even more common in saliva. Reconstructed draft genomes of these two species from metagenomic samples in which they were abundant were combined with modern public genomes to provide a detailed overview of their core genomic diversity.

## 1. Introduction

Multiple research areas have undergone revolutionary changes in the last 10 years due to broad accessibility to high throughput DNA sequencing at reduced costs. These include the evolutionary biology of microbial pathogens based on metagenomic sequencing. Studies on *Mycobacterium tuberculosis* [6,7], *Mycobacterium leprae* [8,9], *Yersinia pestis* [2,10–14] and *Salmonella enterica* [4,15,16] have yielded important insights into the history of infectious diseases by combining modern and historical genomes. In principle, the same approach might also help to elucidate the evolutionary history of both commensal and pathogenic taxa within the human oral microbiome. Periodontitis and dental caries have likely afflicted humans since their origins [17–20]. They may now be amenable to population genetic analyses because a landmark publication by Adler *et al.* in 2013 [21] demonstrated that dental calculus (calcified dental plaque) from the teeth of skeletons that were up to 7500 years old could contain relatively well preserved ancient bacterial DNA. That publication was based on 16S rRNA sequences, which are not informative about intra-species genetic diversity. However, subsequent shotgun sequencing from modern and ancient dental calculus [22–24] has demonstrated that it should be possible to reconstruct genomic sequences that span millennia of human history from multiple individual species within the oral microbiome.

Reconstructing evolutionary history from the oral microbiome faces numerous technical challenges. Our understanding of the historical evolutionary biology of bacterial pathogens benefitted greatly from existing frameworks for the modern population genomic structure of those bacteria [25–27]. However, extensive bacterial population genetic analyses are largely lacking for the modern oral microbiome. The existing literature largely focuses on taxonomic binning into a traditional subset of 40 cultivatable species from periodontitis [28], whose sub-species population structure have not yet been adequately addressed at the genomic level. Instead, most analyses have focused on the “oral complexes”, which consist of groups of multiple species whose co-occurrence is statistically associated with periodontitis [5].

A second barrier to reconstructing evolution history are the limits of the currently existing bioinformatic tools. The genetic diversity of metagenomic sequences is usually classified by binning the microbial sequence reads into taxonomic units. Taxonomic assignments can be performed by the *de novo* assembly of metagenomic reads into MAGs (metagenomic assembled genomes) [29,30], or by assigning individual sequence reads to existing reference genomes. However most current metagenomic classifiers rely on the public genomes in NCBI, whose composition is subject to an extreme sample bias and which represents a preponderance of genomes from pathogenic bacteria [31]. Furthermore, shotgun metagenomes often include DNA from environmental sources, which include multiple micro-organisms that have never been cultivated, and may belong to unknown or poorly classified microbial taxa whose abundance is not reflected by existing databases. Recent evaluations have also demonstrated that current taxonomic classifiers either lack sufficient sensitivity for species-level assignments, or suffer from false positives, and that they overestimate the number of species in the metagenome [31–33]. Both tendencies are especially problematic for the identification of microbial species which are only present at low-abundance, e. g. detecting pathogens in ancient metagenomic samples.

Over the last few years we have developed a series of tools which can facilitate comparative metagenomics of modern and ancient samples. SPARSE, a novel taxonomic classifier for short read sequences in metagenome, was designed to provide accurate taxonomic assignments of metagenomic reads [1]. SPARSE accounts for the existing bias in reference databases [31,34] by sorting all complete genomes of Bacteria, Archaea, Viruses and Protozoa in RefSeq into sequence similarity-based hierarchical clusters with a cut-off of 99% average nucleotide identity (ANI99%). It subsequently extracts a representative subset from those clusters, consisting of one genome per ANI95% cluster because ANI95% is a common cutoff for individual bacterial species [35,36]. SPARSE then assigns metagenomic sequence reads to these clusters by using Minimap2 [37]. However, such alignments are likely to be inaccurate when they are widely dispersed across multiple ANI95% clusters because such wide dispersion reflects either ultra-conserved elements of uncertain specificity or a high probability of homoplasies due to horizontal gene transfer. SPARSE therefore reduces such unreliable alignments by negative weighting of widely dispersed sequences reads. The remaining metagenomic reads are then assigned to unique species-level clusters on the basis of a probabilistic model, and labelled according to the taxonomic labels and pathogenic potential of the genomes within those clusters. Our methodological comparisons demonstrated that SPARSE has greater precision and sensitivity with simulated metagenomic data than 10 other taxonomic classifiers, and yielded more correct identifications of pathogen reads within metagenomes of ancient DNA than five other methods [1]. SPARSE is also suitable for classifying reads from metagenomes from modern samples, and can extract reads from any ANI95% taxon of interest.

SPARSE assigns sequence reads to taxa, but does not create genomic assemblies from the selected metagenomic reads. That task is performed by EToKi, a stand-alone package of useful pipelines that are used by EnteroBase [2] for manipulations of 100,000s of microbial genomes. EToKi is used to merge overlapping paired-end reads, remove low quality bases and trim adapter sequences. It then excludes sequence reads with greater sequence similarities to genomes from a related but distinct out-group than to an in-group of genomes from the target taxon of interest. EToKi then masks all nucleotides in an appropriate reference genome, and creates a pseudo-MAG by unmasking nucleotides with sufficient coverage among the reads that have passed the in-group/out-group comparisons. Finally, EToKi can create a SNP matrix from pseudo-MAGs plus additional draft genomes, and generate a Maximum-Likelihood phylogeny (RAxML 8.2 [38]), which can be visualized together with its metadata in GrapeTree [3].

Here we demonstrate the power of this combination of pipelines by examination of the metagenomic diversity of the human oral microbiome from a large number of historical and modern samples from diverse geographic sources. We address the question of which microbial taxa are uniformly present in human saliva, dental plaque and dental calculus, and which are specific to individual niches. We test the associations of oral taxa within the traditional oral complexes, and conclude that their very existence needs re-examination.

Finally, we examine the population genomic structures of *Streptococcus mutans* and *Streptococcus sobrinus*, which are associated with dental caries in some human populations [39–41].

## 2. Results

### (a) SPARSE analysis of oral metagenomes

We identified 17 public archives containing 1,016 sets of metagenomic sequences (table 1) from 791 oral samples from a variety of global sources which had been obtained from modern human saliva, modern human dental plaque or historical dental calculus (electronic supplementary material, table S1). Individual sequence reads from those metagenomes were assigned to taxa with SPARSE. The assignments were made according to an upgraded database of 20,054 genomes of Bacteria, Archaea or Viruses, one genome per ANI95% cluster among 101,680 genomes in the NCBI RefSeq databases in May 2018. Seven metagenomes (ancient dental calculus: 5; modern saliva: 2) lacked bacterial reads from the oral microbiome (electronic supplementary material, table S2). These seven metagenomes were ignored for further analyses, leaving assignments to 1,591 microbial taxa from 1,009 metagenomes (784 samples) (table 2). Table S3 in electronic supplementary material reports the percentage assignment of the reads in each sample to each of the 1,591 taxa, except for assignments with a sequence read frequency of <0.0001%, which are reported as 0%. Table S3 includes a column identifying assignments to the oral microbial complexes defined by Socransky *et al.* [5]. SPARSE also identified 152 samples containing Archaea from four species, 214 samples containing at least one of four human viruses and 146 samples containing at least one of 12 bacteriophages (table 3). This dataset may represent the currently broadest sample of the oral microbiome from global sources and over time.

**Table 1.** Sources of metagenomic reads.

| Archive | Accession | Sets of short reads | Number of samples | Source | Institute | Citation |
| --- | --- | --- | --- | --- | --- | --- |
| 1 | PRJNA445215 | 62 | 48 | ancient calculus | Max Planck Institute for the Science of Human History | [57] |
| 2 | PRJEB30331, PRJNA454196 | 45 | 44 | ancient calculus | University of Oxford | [24] |
| 3 | PRJNA216965 | 9 | 2 | ancient calculus | University of Oklahoma | [22] |
| 4 | PRJNA383868 | 87 | 87 | plaque | J. Craig Venter Institute | [58] |
| 5 | PRJNA255922 | 48 | 48 | plaque | University of California, Los Angeles | [59] |
| 6 | PRJNA78025 | 7 | 4 | plaque | University of Maryland | [60] |
| 7 | PRJNA289925 | 1 | 1 | plaque | University of Washington | [61] |
| 8 | PRJEB6997 | 298 | 298 | plaque & saliva | BGI | [50] |
| 9 | PRJNA230363 | 12 | 12 | plaque & saliva | Chinese Academy of Sciences | [62] |
| 10 | PRJEB24090 | 61 | 61 | saliva | University of California San Diego | [63] |
| 11 | PRJNA380727 | 56 | 55 | saliva | Peking University School of Stomatology |  |
| 12 | PRJNA396840 | 30 | 30 | saliva | University of Copenhagen | [64] |
| 13 | PRJEB14383 | 28 | 28 | saliva | University College London | [65] |
| 14 | PRJDB4115 | 26 | 26 | saliva | University of Tokyo | [66] |
| 15 | PRJNA217052 | 217 | 18 | saliva | Broad Institute | [67] |
| 16 | PRJNA188481 | 8 | 8 | saliva | Broad Institute | [68] |
| 17 | http://dx.doi.org/10.4225/55/584775546a409 | 21 | 21 | ancient calculus | OAGR, University of Adelaide | [69] |

Note: Ancient calculus refers to ancient dental calculus from historical samples. Plaque and saliva refer to modern dental plaque and saliva.
Sets of short reads were downloaded from GenBank except for Archive 17, which was downloaded from the Online Ancient Genome Repository.
Seven metagenomes (electronic supplementary material, table S2), (Archive 11:2; Archive 17: 5) were excluded from further analyses because they contained too few reads from common microbial taxa in the oral microbiome.

**Table 2.** Sources of genomes from cultivated bacteria and metagenomic samples.

| Category | Sub-category | Number |
| --- | --- | --- |
| Bacterial genomes |  | 262 |
|  | *S. mutans* | 195 |
|  | *S. sobrinus* | 50 |
|  | others | 17 |
| Metagenome source |  | 784 |
|  | Ancient dental calculus | 110 |
|  | Modern plaque | 287 |
|  | Modern saliva | 387 |
| Metagenome size | (nucleotides) |  |
|  | 0-2GB | 343 |
|  | 2-4GB | 129 |
|  | 4-6GB | 162 |
|  | 6-8GB | 93 |
|  | 8-10GB | 45 |
|  | >10GB | 12 |
| Country |  |  |
|  | **Asia** | 442 |
|  | China | 375 |
|  | Japan | 32 |
|  | Philippines | 28 |
|  | Others | 7 |
|  | **North America** | 159 |
|  | U.S.A. | 157 |
|  | Guadeloupe | 2 |
|  | **Europe** | 166 |
|  | U.K. | 75 |
|  | Ireland | 36 |
|  | Denmark | 31 |
|  | Others | 24 |
|  | **Oceania** | 111 |
|  | Australia | 92 |
|  | Fiji | 18 |
|  | Papua New Guinea | 1 |
|  | **Africa** | 9 |
|  | South Africa | 6 |
|  | Sudan | 2 |
|  | Sierra Leone | 1 |

Additional details can be found in electronic supplementary material, table S1.

**Table 3.** Detailed summary of Archaea and Viruses in all 786 samples.

| Taxonomy | No. ancient samples (110) | Percent of reads | No. plaque (287) | Percent of reads | No. saliva (387) | Percent of reads |
| --- | --- | --- | --- | --- | --- | --- |
| **Host (Human)** | **110** | **0.32** | **243** | **9.12** | **335** | **7.05** |
| **Archaea (4)** | **81** | **1.78** | **26** | **2E-4** | **45** | **1E-4** |
| *Methanobrevibacter oralis* | 79 | 1.76 | 26 | 2E-4 | 43 | 1E-4 |
| *Methanobrevibacter smithii* | 1 | 3E-5 |  |  | 2 | 2E-6 |
| *Candidatus Nitrosoarchaeum koreensis* | 1 | 1E-5 |  |  | 0 |  |
| *Thermoplasmatales archaeon BRNA1* | 1 | 7E-6 |  |  | 0 |  |
| **Human viruses (4)** | **0** |  | **25** | **3E-4** | **189** | **4E-3** |
| *Human betaherpesvirus 7* | 0 |  | 8 | 9E-6 | 150 | 6E-4 |
| *Human gammaherpesvirus 4* | 0 |  | 16 | 3E-4 | 86 | 3E-3 |
| *Human alphaherpesvirus 1* | 0 |  | 1 | 5E-6 | 9 | 8E-5 |
| *Human betaherpesvirus 6B* | 0 |  | 0 |  | 7 | 2E-5 |
| **Bacteriophages (12)** | **3** | **1E-5** | **26** | **3E-5** | **117** | **2E-4** |
| *Streptococcus EJ-1* | 0 |  | 14 | 1E-5 | 56 | 8E-5 |
| *Streptococcus SM1* | 2 | 5E-6 | 11 | 1E-5 | 41 | 3E-5 |
| *Streptococcus SpSL1* | 0 |  | 0 |  | 9 | 2E-5 |
| *Streptococcus Dp-1* | 0 |  | 0 |  | 7 | 2E-5 |
| *Streptococcus DT1* | 0 |  | 0 |  | 7 | 2E-5 |
| *Streptococcus PH10* | 1 | 6E-6 | 2 | 3E-6 | 7 | 6E-6 |
| *Klebsiella KP15* | 0 |  | 0 |  | 6 | 6E-6 |
| *Lactococcus r1t* | 0 |  | 0 |  | 6 | 4E-6 |
| *Streptococcus YMC-2011* | 0 |  | 0 |  | 4 | 1E-5 |
| *Propionibacterium PHL060L00* | 0 |  | 0 |  | 2 | 2E-6 |
| *Propionibacterium PHL179* | 0 |  | 0 |  | 1 | 2E-6 |
| *Propionibacterium PAD20* | 0 |  | 0 |  | 1 | 2E-6 |

No. refers to the numbers of samples after combining metagenomes from a common sample.
Percentage of reads refers to the percentage of all reads attributed to a taxon in all the metagenomes from that sample.

### (b) Comparisons of microbiomes from saliva, plaque and historical dental calculus

We tested whether individual oral taxa were particularly enriched or depleted according to source with multiple quantitative approaches, including UMAP (Uniform Manifold Approximation and Projection), principal component analysis (PCA), and hierarchical clustering.

UMAP is a recently described, high performance algorithm for dimensional reduction of diversity within large amounts of data by non-linear multidimensional clustering [42]. A UMAP plot of the taxon abundances in each sample showed three clusters (figure 1A). The three clusters are totally discrete (electronic supplementary material, figure S1A) according to a machine learning approach, optimal k-mean clustering of the first three components from the UMAP analysis). With minor exceptions, the three UMAP clusters were also predominantly associated with source, with one cluster for taxa from modern saliva, a second one for taxa from modern dental calculus and the third for taxa from ancient dental calculus (figure 1A). Similar results were obtained with a classical principal component analysis (PCA), except that the clusters were not as clearly distinguished as with UMAP, and the proportion of exceptions was greater (electronic supplementary material, figure S1B). The assignments of source affiliations to cluster were also largely consistent between UMAP and PCA, with occasional exceptions (electronic supplementary material, figure S1C).

**Figure 1.**
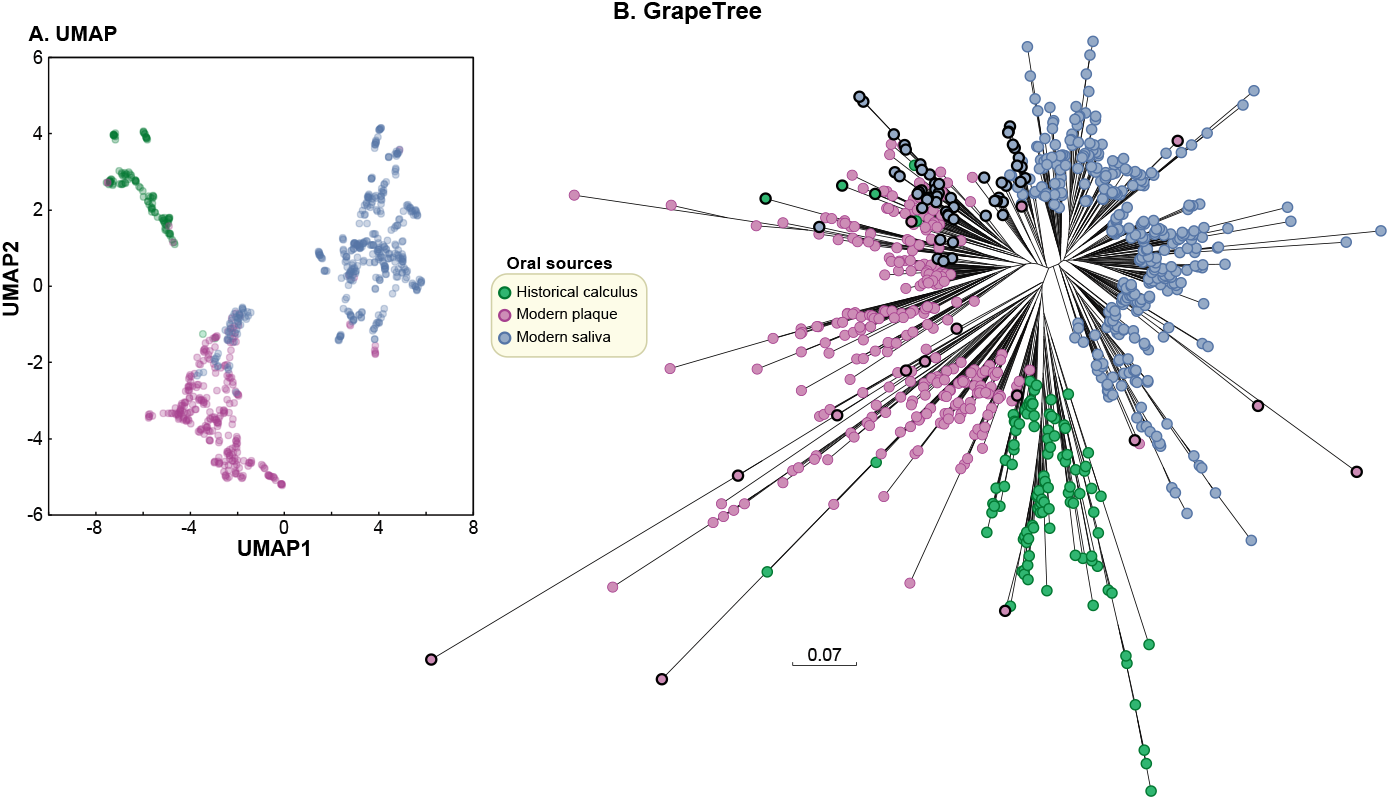
Source specificity of the percentage of species composition in 784 oral metagenomes according to SPARSE. (A) X-Y plot of the first three components from a UMAP (Uniform Manifold Approximation and Projection) [42] dimensional reduction of taxon abundances. (B) Neighbour-joining (FastMe2; [70]) hierarchical clustering based on the Euclidean distances between pairs of metagenomes. Euclidean p-distances were calculated between each pair as the square root of the sum of the squared pairwise differences in the percentage of reads assigned by SPARSE to each microbial taxon. Nodes whose cluster location was inconsistent with the UMAP clustering in part A are highlighted with black perimeters. Tree visualization: GrapeTree [3].

For the third approach, we calculated the Euclidean p-distances between each pair of samples, and subjected them to hierarchical clustering by the neighbor-joining algorithm with the results shown in figure 1B. Hierarchical clustering also largely separated the samples by source with only few exceptions. Samples from modern saliva formed one large cluster. Samples from modern dental plaque formed two related but discrete sub-clusters, one of which included a sub-sub cluster of samples from historical dental calculus. These clusters also largely corresponded to the clusters found by k-mean clustering of UMAP data. Thus, three primary and distinct clusters were consistently identified by three independent methods from the quantitative numbers of reads in individual microbial taxa. The three clusters were largely source-specific for modern saliva, modern plaque and historical dental calculus. This finding predicts that the microbiomes from these three sources contain source-specific taxa.

### (c) Source-specific taxa

We attempted to identify the most important bacterial taxa for the observed clustering by sample source with a second, powerful machine learning approach. A supervised Support Vector Machine (SVM) [43] classification was used to identify the most optimal of 300 SVM model variants, and the 40 most discriminating ANI95% taxa according to that model are shown in figure 2, together with mini-histograms that summarize the relative abundance of sequences by source. As predicted from the discrete clustering described above, multiple taxa were dramatically more prominent in samples from one source than from either of the two other sources. The results also show that the most prominent sample source varied with the taxon (figure 2).

**Figure 2.**
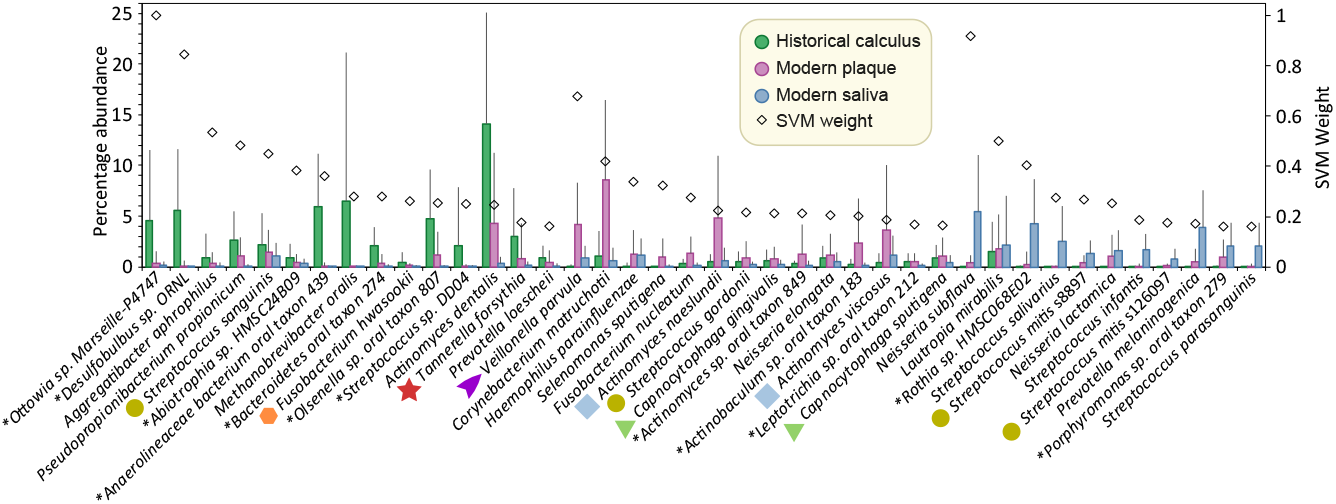
Average percentage abundance (left axis) of bacterial species by source for the 40 most discriminating species according to Support Vector Machine analysis. The relative abundances for each of the three sources are indicated by mini-histograms for each species; error bars indicate standard deviations. Species are sorted in descending order by predominant source and then by SVM weight (squared coefficient) in the optimal model. Species belonging to oral complexes are indicated by oral-complex-specific shapes and colours. Key legend: Source colours used in the mini-histograms and symbol for SVM weight. *species designations assigned by RefSeq to single genomes which have not (yet) been confirmed by taxonomists. *S. mitis* is separated into multiple ANI95% clusters, two of which (s8897; s126097 [electronic supplementary material, table S3]) are among the predominant taxa associated with saliva.

Eleven of the 40 most discriminatory taxa belonged to the oral complexes that are associated with periodontitis according to Socransky *et al.* [5]. Seven species from oral complexes *(Veillonella parvula, Fusobacterium nucleatum, Capnocytophaga gingivalis, Streptococcus gordonii, Actinomyces naeslundii, Actinomyces viscosus,* and *Capnocytophaga sputigena)* were most abundant in modern plaque and two other species *(Streptococcus sanguinis, Tannerella forsythia)* were most abundant in historical dental calculus. The yellow complex includes *Streptococcus mitis,* which encompasses over 50 distinct ANI95% clusters [44]. Two of these ANI95% clusters, designated *S. mitis* s8897 (ANI95% cluster in electronic supplementary material, table S3; MG_43 in [44]) and *S. mitis* s126097 (MG_56) were included among the 40 most discriminatory taxa, and each of them was more frequent in saliva than in dental plaque or dental calculus.

Seventeen other taxa that were assigned to an oral complex by Socransky *et al.* [5] are not included in figure 2 because they were not among the 40 most discriminatory taxa. We therefore examined the relative abundances of all 28 taxa from oral complexes in greater detail (figure 3). Three of the four taxa in the Blue and Purple Complexes are very abundant in oral metagenomes, and all four are preferentially found in modern plaque. However, the other oral complexes are heterogeneous in their patterns of relative abundances. For example, within the Red complex, both *T. forsythia* and *Treponema denticola* were most frequently found in historical dental calculus but *Porphyromonas gingivalis* is most frequent in modern plaque, and is generally much less abundant. Similar intra-complex discrepancies were found for the Orange, Yellow, and Green Complexes. These inconsistent frequencies by source raise questions about the consistency of the compositions of those complexes in individual samples

**Figure 3.**
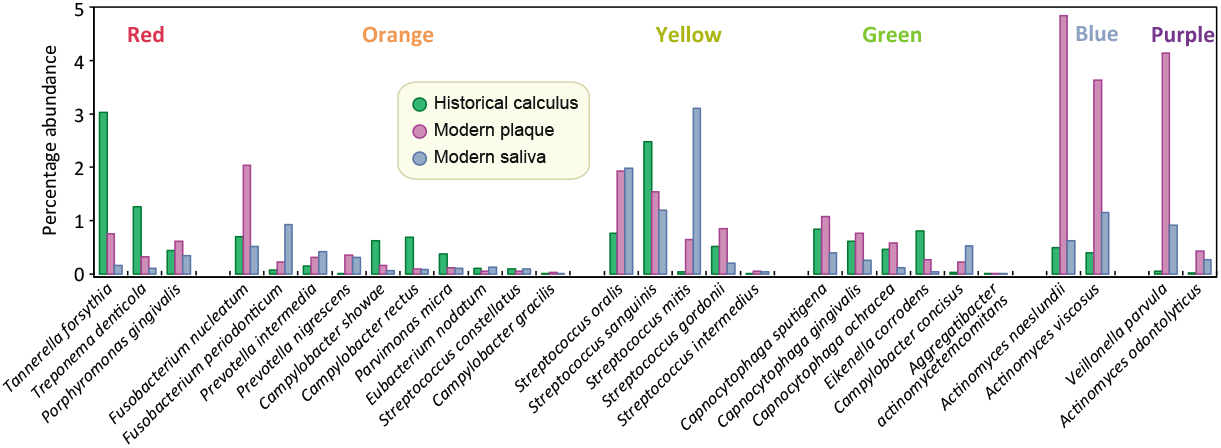
Average percentage abundances in 784 metagenomes by oral source (key legend) of 28 species from six oral complexes described by Socransky *et al.* [5]. The oral sources are indicated by three mini-histogram bars for each species. Species are ordered from left to right by oral complex, whose colours designation is indicated at the top. Within each oral complex, the species order is by decreasing total abundance.

### (d) Existence of “oral complexes”?

Socransky *et al.* [5] initially treated the oral complexes as a hypothesis. However, they have now attained the status of accepted wisdom, and even play a prominent role in routine laboratory investigations and treatment of periodontitis. The oral complexes included 28 cultivated bacterial species, whose presence or absence was determined by DNA hybridization against a small number of probes. This technology is now outdated; the number of known oral taxa has increased dramatically; and the data presented here are for relative abundance rather than presence or absence. However even after weighting for genome size, we do not find a close correspondence between the frequencies of cells in sub-gingival dental plaque measured by Socransky [28] and the results presented here (Supplementary Text). We therefore re-examined the strengths of association with the oral complexes from the data presented here according to similar criteria and similar methods as those used in the Socransky *et al.* 1998 publication [5].

The original assignments to the oral complexes depended strongly on results from hierarchical clustering of the pairwise concordance between species for presence or absence in individual samples. The tree in figure 4 shows neighbor-joining clustering of the common microbial taxa in our dataset by the similarities of their abundances over all samples in our dataset according to SPARSE. This tree contradicts the original composition of the oral complexes: the four areas of the tree where oral complex taxa are clustered each contain representatives from multiple complexes, and none of those four clusters corresponds to the original compositions proposed by Socransky *et al.* [5].

**Figure 4.**
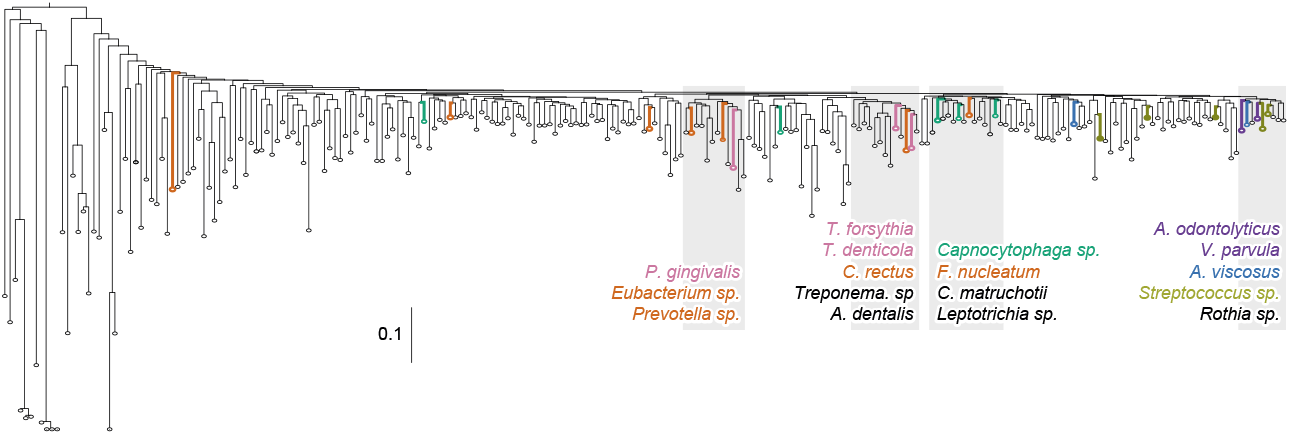
Neighbour-joining (FastMe2; [70]) hierarchical clustering based on the Euclidean distances between pairs of 245 microbial species whose percentage abundance was >2% in at least one metagenome. Members of the six oral complexes [5] are highlighted by coloured species names, whose colours indicate their oral complex membership. These species do not cluster by oral complex, but by other unnamed groupings, four or which are highlighted in gray. An expanded version of the same tree including all species labels is available in electronic supplementary material, figure S2. Branch length distance scale bar is next to the distance of 0.1.

It seemed possible that the discrepancies between figure 4 and the original compositions of the oral complexes might reflect the fact that this study identified many additional taxa, some of which were as common as those used to define the oral complexes (Supplementary Text). We therefore performed cluster analyses of our current data for the original set of 31 cultivatable bacterial species examined by Socransky *et al.* [5]. We compared the neighborjoining algorithm used here with the less powerful, agglomerative clustering method (UPGMA, Unweighted Pair Group Method with Arithmetic Mean) that had been used by Socransky *et al.* We also compared the abundances across all samples with abundances in plaque, which was the primary source for bacteria tested by Socransky *et al.* The results (electronic supplementary material, figure S3) show dramatic inconsistencies between independent trees in regard to the clustering of the oral complex bacteria. For example, *T. forsythia, T. denticola* and *P. gingivalis* of the Red Complex cluster together (and also with *C. rectus)* in electronic supplementary material, figures S3A,C,F,G. However, *T. denticola* and *T. forsythia* are separated from *P. gingivalis* in the four other graphs in electronic supplementary material, figure S3. And none of the three cluster together with each other in electronic supplementary material, figure S3E. Similar, or even greater, discrepancies are visible for the other oral complexes in electronic supplementary material, figure S3. Inconsistencies in clustering patterns across minor differences in sampling and clustering algorithms raise severe doubts about the very existence of the oral complexes as defined by Socransky *et al.* [5].

### (e) Numbers of taxa per source

The rarefaction curves in figure 5A provide a breakdown of taxa by sample source as additional samples are tested. SPARSE detected 1591 microbial taxa over all 784 metagenomic samples: 1,389 from modern saliva; 842 from modern plaque and 696 from historical calculus. These estimates will increase as additional samples are added, but at increasingly slower rates because the rarefaction curves seem to be reaching a plateau, except for historical dental calculus where the fewest samples have been evaluated until now.

**Figure 5.**
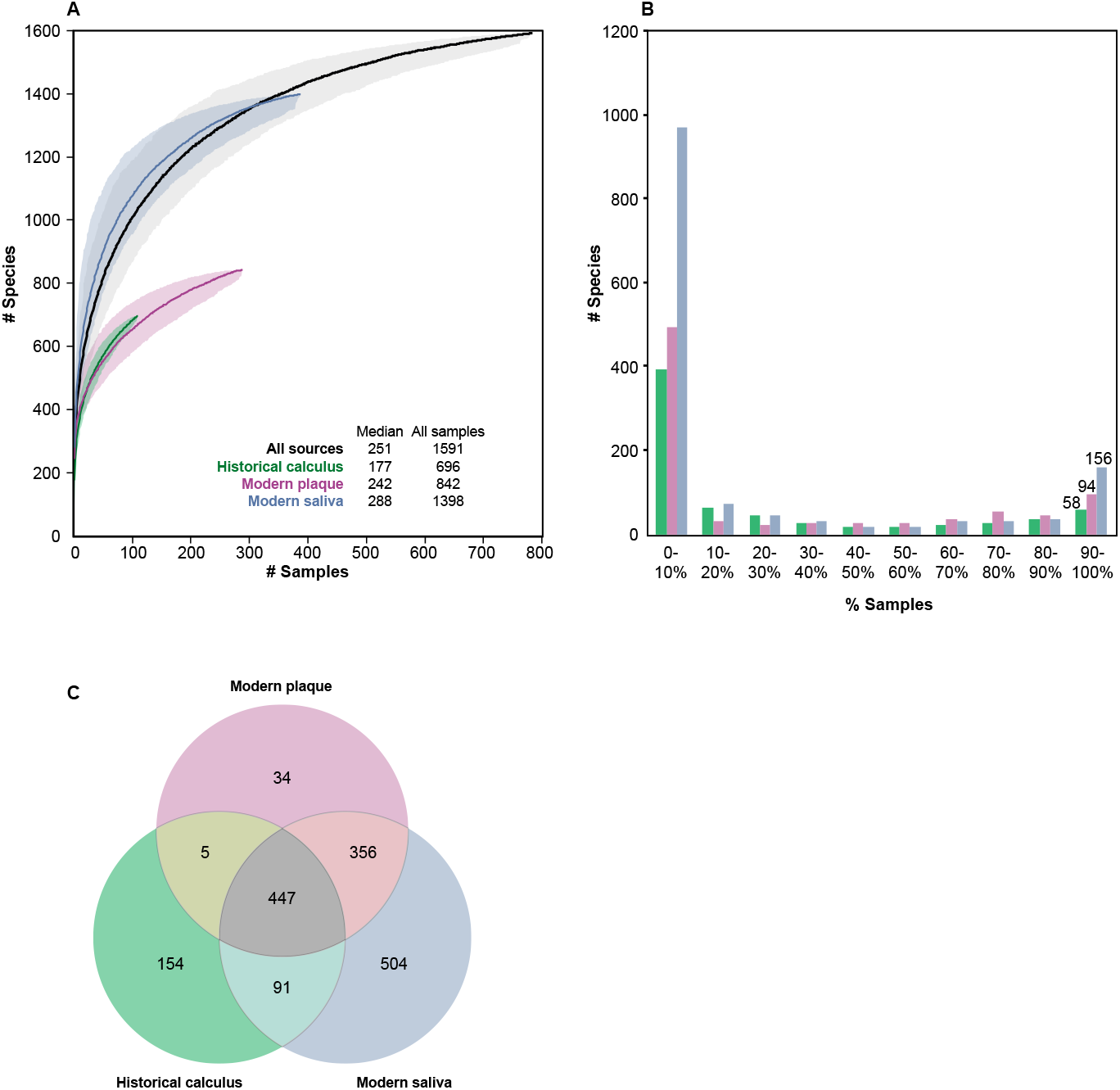
Numbers of microbial taxa by source. A). Rarefaction curves of numbers of species by source, with 95% confidence estimates (shadow). Inset data indicates median numbers of species per sample by source, as well as the total numbers for all sources. Rarefactions were performed with the program script called SPARSE_curve.py, using 1000 randomized permutations of the order of samples. B). Binned histograms of number of species by percentage of samples. The data for this plot was also calculated with SPARSE_curve.py. C) Venn diagram of overlapping presence of taxa (≥0.0001% abundance) for the three oral sources.

The median numbers of taxa per sample range from 177 (historical dental calculus) to 288 (modern saliva), and were much smaller than the total numbers. These median values reflect a bimodal distribution for numbers of taxa per sample (figure 5B), wherein a few samples had jackpots of large numbers of taxa but all other samples had only few.

The analyses described above focused on differences in taxon composition by source. However, the Venn diagram in figure 5C shows that 447 taxa were common to all three sources, even if their relative abundances varied. Modern plaque yielded only 34 taxa which were not found in either historical dental calculus or modern saliva. More source-specific taxa were found in historical dental calculus, which may possibly reflect some contamination with environmental material. Alternatively, some taxa may be absent in modern dental plaque because historical lineages have become extinct [4]. Saliva yielded 504 unique taxa, some of which might be transient, and do not persist long enough to be incorporated into plaque.

### (f) Population genomics of organisms associated with dental caries

The microbiome associated with early stages of dental caries is an unresolved topic that remains under active investigation [40,45–47]. However, it is generally accepted that *Streptococcus mutans* and *Streptococcus sobrinus* are routinely associated with caries [48]. Our data confirm that reads belonging to these two taxa are abundant in modern dental plaque, and also show that they are even more abundant in modern saliva (figure 6A,C). However, there was no significant correlation between the relative frequencies of these species across multiple metagenomes (electronic supplementary material, figure S9). Prior analyses based on 16S RNA OTUs indicated that *S. mutans* was extremely rare in historical dental calculus, and argued that this increase was caused by the introduction of high levels of sugar to human diets in industrialized societies in the last 200 years [21]. Our data show that *S. sobrinus* was undetectable in historical samples (frequency of <0.0001% of reads or <10 reads per metagenome) (figure 6C). *S. mutans* was also undetectable in most of these samples, but up to 0.04% of all reads in 10 historical samples spanning the last 1500 years were assigned to *S. mutans* (figure 6A), in accordance with archaeological findings that dental caries has been common in multiple eras over the last 10,000 years [17]. The few reads from historical samples that were assigned to *S. mutans* showed increased deamination at their 5’-ends when tested by MapDamage2 [49] (electronic supplementary material, figure S4), confirming that they were truly from ancient DNA.

**Figure 6.**
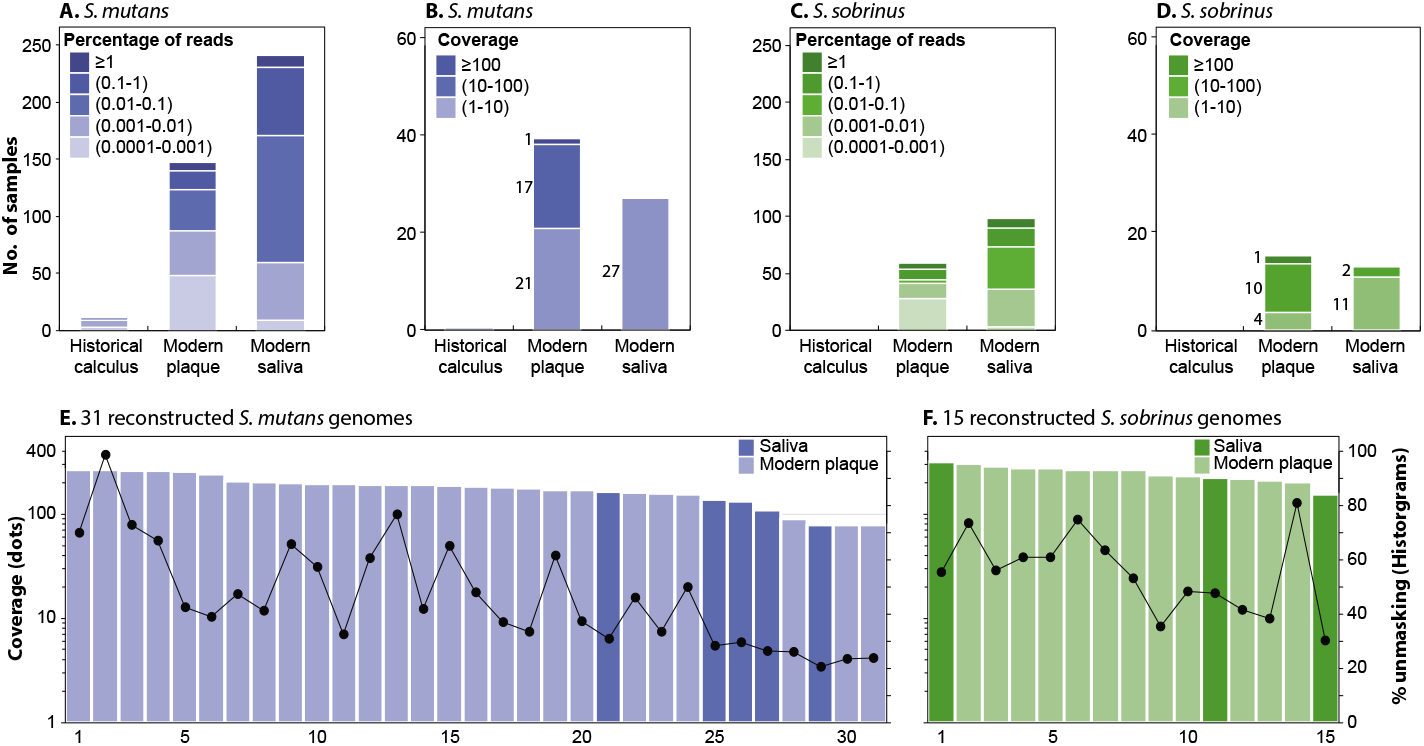
Reconstruction of pseudo-MAGs (metagenomic assembled genomes) of *S. mutans* and *S. sobrinus* from oral metagenomes. (A, C) Numbers of oral samples by source binned by the percentage of reads specific to *S. mutans* (A) and *S. sobrinus* (C). (B, D) Numbers of oral samples by source with an average coverage of at least 1x. The data are binned by the predicted read coverage against a reference genome of *S. mutans* (UA159) (B) and *S. sobrinus* (NCTC12279) (D). (E, F) Read coverage (Dots; left) and percentage of the reference genome that was unmasked (≥3 reads; ≥70% consistency) (Histogram; right) in *S. mutans* (E) and *S. sobrinus* (F). Ordered by decreasing coverage.

We exploited the high frequency of sequence reads from these two *Streptococcus* species in modern dental plaque and saliva to illustrate how SPARSE and EToKi can be used to extract pseudo-MAGs from metagenomic sequence reads, and combine them with genomes sequenced from cultivated bacteria (Methods). These procedures resulted in a total of 31 pseudo-MAGs for *S. mutans* and 15 pseudo-MAGs for *S. sobrinus* in which over 70% of the reference genome had been unmasked (figures 6E,F, electronic supplementary material, table S6). Most of these pseudo-MAGs were from Chinese samples [50]. The pseudo-MAGs were combined with genomes from cultivated bacteria of the same species from Brazil, the U.S. and the U.K. as well as other countries (table 2) and Maximum Likelihood (ML) phylogenies of non-repetitive SNPs (figure 7) were created with EToKi (Methods).

**Figure 7.**
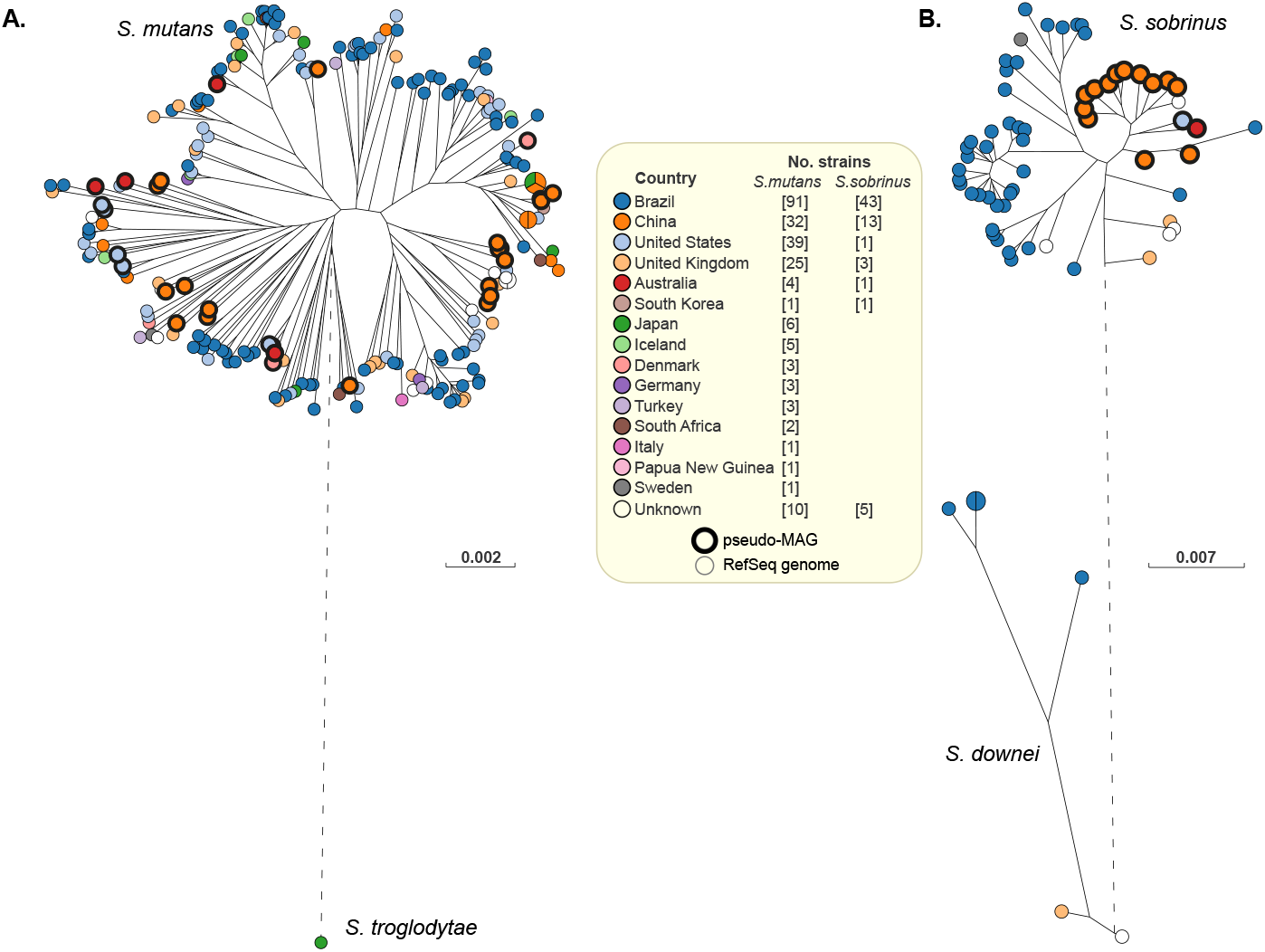
Maximum Likelihood phylogenies of *S. mutans* and *S. sobrinus* genomes. (A) A RaxML [38] tree of 226 genomes of *S. mutans* (RefSeq: 195; pseudo-MAGs: 31) plus one genome of *S. troglodytae* as an outgroup. The tree was based on 181,321 non-repetitive SNPs in 1.73 Mb. (B) A RaxML tree of 61 genomes of *S. sobrinus* (RefSeq: 46; pseudo-MAGs: 15) plus six *S. downei* genomes as an outgroup. The tree was based on 160,863 non-repetitive SNPs in 1.13 Mb. Pseudo-MAGs are highlighted by thick black perimeters. Visualisation with GrapeTree [3]. Branches with a genetic distance of >0.1 were shortened for clarity, and are shown as dashed lines. Legend: Numbers of strains by country of origin for both trees.

The ML phylogenies of the two species showed interesting differences. All 13 Chinese pseudo-MAGs clustered together within the *S. sobrinus* ML tree (figure 7B), whereas almost all the other 44 bacterial genomes from Brazil and elsewhere clustered distantly. In contrast, in the *S. mutans* tree (figure 7A), 20 Chinese pseudo-MAGs did not show any obvious phylogeographic specificities, and were inter-dispersed among 196 bacterial genomes from multiple geographic locations. Similar conclusions about a lack of phylogeographic specificity were previously reached by Cornejo *et al.* [51] on a subset of 57 of these *S. mutans* genomes.

## 3. Discussion

Several years ago, we accidentally became interested in comparing historical and modern genomes reconstructed from metagenomic short read sequences with draft genomes assembled from high throughput sequencing of cultivated bacteria. Our initial efforts involved the deployment of individual bioinformatic tools, comparisons of multiple publicly available algorithms, and compilation of draft genomes from publicly available sequence read archives of short read sequences [7]. In parallel, we were also involved in developing EnteroBase, a compendium of 100,000s of draft genome assemblies from multiple genera that can cause enteric diseases in humans, including *Salmonella* [2,27]. These two projects were synergistic for elucidating the evolutionary history of *Salmonella enterica* based on metagenomic sequences from 800-year old bones, teeth and dental calculus [4]. In that case, sequence reads from *S. enterica* were found in teeth and bone, but not in dental calculus. Our attempts to examine further samples of dental calculus quickly demonstrated that optimized pipelines were needed because manual analyses were too time-intensive. However, none of the existing tools were both reliable and sufficiently sensitive for assigning sequence reads from historical metagenomes to the tree of microbial life. We therefore took a step back, and developed SPARSE [1] to satisfy our requirements. SPARSE replaces the current reference databases, which are strongly biased to multiple, closely related genomes from bacterial pathogens, by a representative subset consisting of one genome per ANI95% hierarchical cluster within RefSeq, and assigns sequence reads to these clusters using a probabilistic model. That model penalizes non-specific mappings of reads, and hence reduces false-positive assignments. SPARSE was more reliable than multiple other taxonomic classifiers, and both more sensitive and more reliable for identifying low numbers of reads from ancient metagenomes than multiple other pipelines [1]. In parallel, we expanded the capacities of EToKi [2], an efficient backend pipeline for genomic manipulations, such that it can accurately identify individual sequence reads sieved through SPARSE that are more similar to an in-group of reference genomes from the target species than to an out-group of genomes from a closely related, but distinct taxon. Those reads are then used to unmask nucleotides in a reference genome and generate a pseudo-MAG for SNP-based maximum likelihood phylogenies. Finally, we developed GrapeTree [3], which facilitates the graphic visualization and manipulation of phylogenetic trees based on large numbers of genomes. Here we demonstrate how to combine all three tools in order to obtain an overview of the microbial flora in samples from human oral saliva, modern dental plaque and historical dental calculus. We also reconstructed genomes of two taxa present at moderate concentrations within the oral microbiome, and compare them with conventional draft genomes. The experimental procedures for processing 1016 metagenomes consisted of running SPARSE in the background for 2 months (~100,000 CPU hours). The pipelines described here permitted all other procedures and evaluations described here to be completed in less than two weeks.

Our traditional understanding of oral ecology is largely based on taxonomic assignments of cultivatable bacteria, often performed by checkboard DNA-DNA hybridization [28]. Currently, 756 species have been cultivated from the human oral cavity and respiratory tract [52]. A subset of 40 are used for checkboard DNA-DNA hybridization [28], of which 28 were used to define the oral complexes that were thought to be of importance for periodontitis [5]. Our comparisons of those data with the results from the metagenomic analyses presented here shows that the frequencies of individual taxa determined by the checkerboard assay were inconsistent with the frequencies determined by our metagenomic analyses (electronic supplementary material, figures S5 and S6). The checkerboard assays also lacked 17 common taxa from dental plaque and dental calculus that were found by metagenomic analyses. These results are not unexpected because our metagenomic analyses included saliva samples as well as ancient dental calculus, and identified 1591 taxa, many of which have not been cultivated. Furthermore, it is now well established that the frequencies of certain supposed members of the oral complexes differ very dramatically with geographical source [53]. However, we had anticipated that we might be able to expand the compositions of the oral complexes to include previously uncultivated organisms. Instead, we were unable to reliably identify their very existence (figure 4) because clustering of taxa was affected by minor changes in choice of samples and choice of clustering algorithm (electronic supplementary material, figure S3). We therefore conclude that the existence and composition of the oral complexes needs independent verification by modern techniques and new samples.

The data presented here provide an unprecedented comparative overview of the relative proportions of the predominant taxa in public available metagenomes from the modern and historical oral microbiome. Figure 2 identifies 15 taxa, which are particularly common in historical calculus, 13 others that are preferentially found in modern dental plaque and 11 that seem to be specific for saliva. These associations with a particular source in the oral cavity might be used to identify currently undefined ecological complexes of oral taxa that share a common niche. However, species-level OTUs are likely to be conglomerates of multiple microbial populations, each of which may inhabit a somewhat different ecology. For some organisms such as *Salmonella* or *Escherichia,* efforts are currently underway to develop hierarchical clustering of such populations in order to categorize their ecological and pathogenic differentiation [2]. A step in this direction for the oral microbiome is the recognition of ANI95% clusters s8897 and s126097, both of which were preferentially found in saliva. A large study of all streptococci [44] identified multiple other ANI95% clusters within *S. mitis* but their preferential location in the oral cavity have not yet been addressed. Indeed, little is yet known about the sub-species population structure of almost all of the taxa identified here.

Our more detailed investigation of *S. mutans* and *S. sobrinus* may represent a forerunner of future studies on sub-species ecological differences within the oral microbiome. *S. mutans* and *S. sobrinus* are commonly associated with dental caries, and may play a causal role in that disease [48]. However, once again these taxa were more common in saliva than in dental plaque (figure 6). We chose *S. mutans* and *S. sobrinus* for more detailed analysis because sufficient reads were found in multiple metagenomes from modern samples to allow the partial reconstruction of multiple genome sequences (pseudo-MAGs). In addition, multiple draft genomes from cultivated bacteria existed in the public domain which were available for genomic comparisons. We were also intrigued by the claim that *S. mutans* was rare in historical plaque [21]. Our data support that claim, and we found only few historical samples of dental calculus that contained any reads of *S. mutans,* and none with *S. sobrinus.* Our data also support prior conclusions of a lack of phylogeographic differentiation within *S. mutans* [51]. However, although the data are still somewhat limited, *S. sobrinus* from China tend to cluster distinctly from genomes from Brazil (figure 7). Distinct clustering might reflect phylogeographical signals but other causes of clustering cannot currently be excluded because the Chinese genomes were pseudo-MAGs reconstructed from metagenomes from dental plaque and saliva while the Brazil genomes were from bacteria cultivated from dental plaque. Additional genomes of *S. sobrinus* from other geographical areas would be needed to determine whether the apparent phylogeographical trends are robust. Such analyses could also be facilitated by creating an EnteroBase for *Streptococcus,* which could be done relatively easily [44] if there were interested curators and sufficient interest in the *Streptococcus* community.

In summary, we illustrate the use of a variety of reliable, high throughput tools for determining microbial diversity within metagenomic data, and for extracting microbial genomes from metagenomes. We illustrate these tools with metagenomes from both modern and historical samples, and release all the data and methods for further use by others.

## 4. Methods

### (a) SPARSE database update

In its original incarnation in August 2017 [1], SPARSE used MASH [54] to assign 101,680 genomes from the NCBI RefSeq database to 28,732 ANI99% clusters of genomes. By May 2018, 21,540 additional genomes had been added to NCBI RefSeq. These were merged into the existing database in the same manner as previously, by merging that genome into an existing ANI99% cluster or by creating a new cluster containing one genome if the ANI to all existing clusters was less than 99%. An ANI99% representative microbial database was generated which contained one representative genome for each of the 32,378 ANI99% clusters containing Bacteria, Archaea or Viruses plus a human reference genome (Genome Reference Consortium Human Build 38) such that reads from human DNA could also be called. All the representative genomes were assigned to a superset of 20,054 ANI95% clusters, and this was used for species assignments and genomic extractions as described [1].

### (b) SPARSE analyses

‘EToKi prepare’ was used to collapse paired-end reads and trim all sequence reads. Subsequent SPARSE analyses were performed on all the metagenomes in table 1 and additional metagenomes in electronic supplementary material, figure S7 as described in the SPARSE manual (https://sparse.readthedocs.io/en/latest/). The first step was ‘SPARSE predict’, which identifies ANI95% groups containing ≥10 specific reads. Subsequently, ‘SPARSE report --low 0.0001’ was used to assign taxon designations to the ANI95% groups, and produce a table of all metagenome results (electronic supplementary material, table S3) which lists distinct taxa for each metagenome that accounted for ≥0.0001% of all its reads. Table S3 also includes the designations of oral complexes and other known pathogens according to a manually curated dictionary. Sequence reads were extracted from the metagenomes for assembling pseudo-MAGs with ‘SPARSE extract’.

For electronic supplementary material, figures S5-S8, the taxonomic assignments were inversely weighted by genome size in order to render them comparable to DNA-DNA Checkerboard data and output from Metaphlan2, which calculate cell counts. To this end, the number of metagenomic reads assigned to each species within a metagenome was divided by the genome size of the SPARSE reference genome for that species. These data were then expressed as a proportion of the summed data for all microbial species within that metagenome.

### (c) Metagenomes lacking reads from the oral microbiome

We tested all metagenomes to identify any that might be grossly contaminated by collating the fifty most abundant microbial species over all metagenomes (electronic supplementary material, table S4A). The percentage of reads in these 50 taxa was summed for each metagenome, and expressed as a percentage of all microbial reads. Seven metagenomes (ancient dental calculus: 5; modern saliva: 2; electronic supplementary material, table S2) were excluded because the percentages of those top oral microbes constituted < 15% of their total microbial reads.

### (d) Dimension reduction of frequencies of reads

Two forms of dimensional reduction of diversity were used to detect source-specific clustering within the SPARSE results. UMAP analysis was performed with its Python implementation [42], using the parameters min_neighbors=5 and min_dist=0.0. PCA was performed using the decomposition.PCA module of the scikit-learn Python library [55]. Optimal k-mean clusters of the first three components from the UMAP analysis were calculated with the sklearn.cluster module of the scikit-learn Python library.

### (e) Ranking of microbial species by their associations with source

Microbial species were ranked by their weighting according to a Support Vector Machine (SVM) classification [43]. A supervised SVM classification of samples was performed using the SVM module of the scikit-learn Python library on the raw SPARSE results (electronic supplementary material, table S3). The SVM classification was performed 300 times on a randomly chosen training set consisting of 60% of all samples with varying penalty hyperparameter C, and scored using 5-fold cross-validation. The model was then tested with the optimal hyper-parameter from all runs on the remaining 40% of samples, and correctly inferred the oral source for >96% of the test samples. The optimal SVM coefficients for each individual species were estimated by training that model once again on all the oral samples. The order of the species in figure 2 consists of the SVM weights (squares of the coefficients; [56]) in descending order. The Python scripts described in sections d and e, as well as their outputs are freely accessible online as Dataset S3 in https://github.com/zheminzhou/OralMicrobiome.

### (f) Genome reconstructions for *Streptococcus mutans* and *Streptococcus sobrinus*

SPARSE identified samples in which the metagenomic sequence reads covered at least 2MB of the reference genome for *S. mutans* (ANI95% cluster s5; 66 samples) or *S. sobrinus* (s3465; 28 samples) (figures 4B,D). The cleaned, species-specific reads generated from these samples as in Methods b were processed with the standalone version of EToKi as described in figure S6 of Zhou *et al.* 2020 [2] and in greater detail in the online manual (https://github.com/zheminzhou/EToKi). EToKi assemble was then used to identify genome-specific reads after specifying a reference genome, an in-group of related genomes, and a related but distinct out-group of other genomes. For *S. mutans* the reference genome was UA159 (accession code GCF_000007465), the in-group was 194 other *S. mutans* genomes in RefSeq (electronic supplementary material, table S5) and the outgroup was 62 genomes from other species in the Mutans *Streptococcus* group according to Zhou and Achtman, 2020 [44]. For *S. sobrinus* the reference genome was NCTC12279 (accession code GCF_900475395), the ingroup was 45 other *S. sobrinus* genomes and the outgroup was 211 genomes from other Mutans streptococci (electronic supplementary material, table S5). The assemble module replaces nucleotides in the reference genome by their calculated SNVs after checking that they are supported by at least 70% of at least 3 metagenomic reads, and that the supporting read frequencies are at least one-third of the average read depth. The resulting pseudo-MAGs are listed in electronic supplementary material, table S6 and are freely accessible online as Datasets S1 and S2 in https://github.com/zheminzhou/OralMicrobiome.

‘EToKi align’ was used to create an alignment of non-repetitive SNPs from 31 *S. mutans* pseudo-MAGs plus all 195 *S. mutans* genomes plus the sole *S. troglodytae* genome in RefSeq (electronic supplementary material, table S5). The alignments spanned 1.73 MB that were shared by ≥ 95% of the genomes, and covered 181,321 core SNPs. Similarly, an alignment of 15 *S. sobrinus* MAGs, 46 draft or complete *S. sobrinus* genomes plus 6 genomes of *Streptococcus downei* from RefSeq spanned 1.16 MB and contained 160,863 core SNPs. These alignments were subjected to Maximum Likelihood phylogeny reconstruction by EToKi phylo. Both ML trees were then visualised with GrapeTree [3].

### (g) DNA damage patterns for ancient *S. mutans* reads

SPARSE assigned low numbers of sequence reads to *S. mutans* in 10 metagenomes from ancient dental calculus (figure 6, electronic supplementary material, table S3). In order to assess their authenticity, these reads were assessed with MapDamage2 [49]for patterns of cytosine deamination that are characteristic of authentic ancient DNA. To this end, all *S. mutans*-specific reads were extracted with SPARSE. They were aligned to the *S. mutans* reference genome UA159 with Minimap2 [37], and reads which were ≥95% identical with the reference genome were used to create BAM alignments. SouthAfr2 contained 11 specific reads according to SPARSE, but only eight survived this filtering step. SouthAfr2 was therefore excluded from further analyses because these were too few reads to provide reliable analyses. The BAM alignments from the remaining nine metagenomes consist of both fully aligned reads (46-72%) and others which were “soft-clipped”, i.e. terminal bases were not aligned to the reference genome. In order to ensure that these soft-clipped reads were also specific, we compared the alignment scores for all reads against UA159 with the alignments scores against the 62 outgroup genomes in Mutans Streptococci (electronic supplementary material, table S5), and found that the scores with UA159 were highest. We also tested the alignment scores against two other *S. mutans* genomes (SA38, [GCF_000339615]; 4VF1 [GCF_000339215]; electronic supplementary material, table S5), but neither yielded higher alignment scores than UA159. The outputs from MapDamage2 show the soft-clipping ends by a yellow line (electronic supplementary material, figures S4A- D).

## Data availability

The pseudo-MAGs reconstructed from metagenomes for *S. mutans* and *S. sobrinus* are freely accessible in tar.gz files containing <u>Datasets_S1</u> and <u>Dataset_S2</u> at https://github.com/zheminzhou/OralMicrobiome, respectively. Python scripts that were used to prepare data for figures 1–5 and S1-S3 are available as <u>Dataset_S3</u> in the same repository. The taxonomic profiling by SPARSE of all 784 metagenomes is available in electronic supplementary material, table S3. Interactive versions of Figure 7 are available at http://enterobase.warwick.ac.uk/a/42277 (figure 7A) and http://enterobase.warwick.ac.uk/a/42279 (figure 7B)

## Authors’ contributions

Z.Z. analysed data and prepared the figures. M.A. and Z.Z. interpreted the results and wrote the manuscript.

