## Supplemental Text for "Metagenomics of the modern and historical human oral microbiome with phylogenetic studies on *Streptococcus mutans* and *Streptococcus sobrinus*"

### Supplementary text

#### (a) Comparisons between SPARSE and DNA-DNA hybridization

Our SPARSE results cast doubt on the existence of the oral complexes which had been proposed in 1998 [5] on the basis of checkerboard DNA-DNA hybridization [28]. Those assays were performed on bacterial DNA which was extracted by boiling sub-gingival dental plaque samples in 0.25M NaOH for five minutes. Bacterial concentrations were measured by probing those extracts with genomic DNA from one representative of each of 40 test species. This approach is potentially error-prone because the efficiency of DNA extraction by boiling may vary with the species, and because the probe DNA includes DNA stretches from the accessory genome whose presence is highly variable within a species. We therefore compared the cell counts of the 40 probe species from checkerboard assay in plaque samples from 776 individuals that had been tested in 2005 [28] with percentage assignments according to SPARSE in 784 oral metagenomes for the same 40 taxa plus other common taxa. These percentage differences were inversely weighted by the reference genome size for each taxon in order to account for differences in numbers of reads with genomic size.

The comparisons revealed large quantitative differences between the two analyses (electronic supplementary material, figure S5). Only 12 of the 40 classical taxa were particularly common (≥ 1%) in metagenomes from saliva, dental plaque or ancient dental calculus. *P. gingivalis* was rare in the metagenomic analyses, and *Campylobacter gracilis, Aggregatibacter actinomycetemcomitans* (aggressive juvenile periodontitis [71]) and *Propionibacterium acnes* (now *Cutibacterium acnes* [72]) were very rare. Sixteen other species that were common within the metagenomes according to SPARSE were not included among the 40 used for checkerboard analyses although all but five of them had already been defined before 1998. The most common of these additional taxa were *Corynebacterium matruchotii* (which plays a primary structural role in dental plaque [73]), *Actinomyces dentalis* (which has been isolated from a root abscess [74]) and *Actinomyces sp.* oral taxon 448. One possible explanation for these discrepancies is that many or most of the modern oral metagenomes were from individuals without periodontal disease whereas the 40 species tested in checkerboard analyses consisted of taxa that were particularly relevant for periodontitis. In order to test this possibility, we subdivided the SPARSE assignments by the status of the plaque donors for periodontitis (electronic supplementary material, figure S6). Figure S6 also include SPARSE results with additional metagenomes recently described by Velsko *et al*. [24] from 20 saliva samples in HMP from the U.S.A. plus 10 samples of modern dental calculus from Spain (see below).

Of the four rare species described above, *P. gingivalis* was indeed found almost exclusively in individuals with periodontal disease (electronic supplementary material, figure S6). Unexpectedly, it was only common in modern plaque, and was extremely rare in modern or ancient dental calculus and saliva. *C. gracilis* and *C. acnes* remained rare in all samples, and *A. actinomycetemcomitans* was only found in saliva. Furthermore, *Prevotella melaninogenica* was the sixth most common taxon in the classical sub-gingival plaque data but was only detected by SPARSE in modern saliva. Interestingly, the most common taxon among the six samples of modern dental calculus from donors with periodontitis [24] was *Actinobaculum* oral taxon 183, which was not included in the checkerboard analyses. Figure S6 demonstrates that the SPARSE data do not reflect the same distributions of taxa as obtained in classical analyses with the checkerboard assay.

We conclude that the checkerboard data do not provide a general overview of the frequencies of taxa that are common in the entire oral microbiome with modern techniques. We also raise the possibility that the checkerboard results might have been distorted by unrecognized technical problems. Alternatively, the discrepancies may also reflect biogeographical differences between the frequencies of different taxa in different countries and continents, as has been noted by others [53].

#### (b) Metaphlan2 *versus* SPARSE

A recent publication by Velsko *et al.* [24] compared metagenomes from dental plaque from 20 healthy individuals in the United States, dental calculus samples from 10 individuals with dental caries in Spain (6 with periodontitis) and 43 samples of 200 year old dental calculus from England of varied status for dental caries and periodontitis. Velsko et al. claimed that the results demonstrated “distinct taxonomic … profiles between plaque, modern calculus, and historic calculus, but not between calculus collected from healthy teeth and periodontal disease-affected teeth”.

Velsko et al. 2019 was published after our SPARSE calculations had been completed for this manuscript. The metagenomes from ancient dental calculus had been included in our analyses, but not the metagenomes from modern dental calculus or modern dental plaque. We describe SPARSE results with those samples here due to a request by one of the reviewers for this manuscript. Our comparison between the SPARSE results and the summaries published by Velsko et al. [24] revealed quantitative discrepancies in the relative frequencies of reads (electronic supplementary material, figure S7A,B), and also showed that SPARSE identified additional taxa that were absent in that publication. We attributed these differences to differential taxonomic assignments between SPARSE and Metaphlan2 [75], a widely cited software package based on a proprietary database of clade-specific markers that had been used by Velsko et al.

Our previous comparisons [1] had shown that Metaphlan2 failed to correctly identify two of three sets of ancient DNAs in low concentration, but we had never previously performed a head to head comparison between SPARSE and Metaphlan2 for more amenable metagenomes, such as those from the modern oral microbiome. We therefore performed a quantitative comparison of our SPARSE results with the assignments of reads from all 73 metagenomic samples according to Velsko *et al*. [24]. Velsko *et al*. assigned reads to 16 genera, which were not directly comparable with the species level assignments by SPARSE. 173 of the other 183 taxa in Velsko *et* al. were subsumed into 164 distinct species by SPARSE at its minimal presence/absence cut-off level of ≥0.02% of all microbial reads. SPARSE also identified reads from 121 other species (electronic supplementary material, figure S7C) at the same cut-off level, most of which are associated with the normal oral microbiome.

The quantitative assignments for numbers of reads assigned by the two pipelines were comparable for 157 taxa that were common to both approaches. However, SPARSE identified dramatically higher or lower proportion of reads than the results in Velso *et al.* for seven other taxa (electronic supplementary material, figure S8A). These differences are large enough to change the overall taxonomic composition of individual metagenomes, as illustrated for sample CS36 (electronic supplementary material, figure S8B). We initially interpreted these differences as reflecting differences between Metaphlan2 and SPARSE, and ran Metaphlan2 over the same data. Metaphlan2 (V2.7.8) assigned a large number of reads in CS36 to *Eubacterium saphenum* and a moderate number to *Fretibacterium fastidiosum*, where SPARSE found only a modest number of reads from *E. saphenum* and none from *F. fastidiosum*. SPARSE also identified numerous reads from the taxa *Peptostreptococcaceae* oral taxon 113, *Anaerolineaceae* oral taxon 439, *Actinomyces Israeli*, and *Actinomyces dentalis* whereas Metaphlan2 found none. However, even these findings did not totally explain the discrepancies with the results in [24], because the supplementary information in that publication contained much higher proportions of *Treponema denticola* and much lower frequencies of *Campylobacter rectus* than Metaphlan2 (2.7.8), SPARSE or MetaPlan2 2.5.0, which was the version that had been used by Velsko *et al.* (electronic supplementary material, figure S8B). In order to clarify the true proportions of these three taxa, we resorted to still a third approach, namely Minimap2 [37], which maps single reads to their position within a complete genome. The results are concordant with those by SPARSE (electronic supplementary material, figure S8C) in that numerous reads were found for *C. rectus,* intermediate numbers for *E. saphenum* and low numbers for *T. denticola.*

We conclude that recent versions of Metaphan2 cannot correctly assign some common taxa within the oral microbiome to a species, and that it overestimates or underestimates the numbers of reads from other common taxa. Due to the additional discrepancies between the Metaphlan results and the Supplementary data published by Velsko *et al*. [24], we are hesitant to compare the SPARSE results presented here with those published by Velsko *et al*. [24], and have otherwise largely ignored them in the main text.
