## Supplementary material for "Metagenomics of the modern and historical human oral microbiome with phylogenetic studies on *Streptococcus mutans* and *Streptococcus sobrinus*": Table S2

Table S2. Oral samples that were excluded from further analysis because <15% of their microbial reads were from the 50 most common oral taxa.

| **Sample** | **Source** | **MBases** | **Center Name** |
| --- | --- | --- | --- |
| EBC2 | dental calculus | 17 | OAGR, University of Adelaide |
| EBC3 | dental calculus | 60 | OAGR, University of Adelaide |
| Spy2 | dental calculus | 288 | OAGR, University of Adelaide |
| Sudan1 | dental calculus | 11 | OAGR, University of Adelaide |
| Sudan2 | dental calculus | 27 | OAGR, University of Adelaide |
| SRS943643 | saliva | 10,490 | BIOLS |
| SRS943644 | saliva | 3,820 | BIOLS |
