## Supplementary material for "Metagenomics of the modern and historical human oral microbiome with phylogenetic studies on *Streptococcus mutans* and *Streptococcus sobrinus*": Figure S1

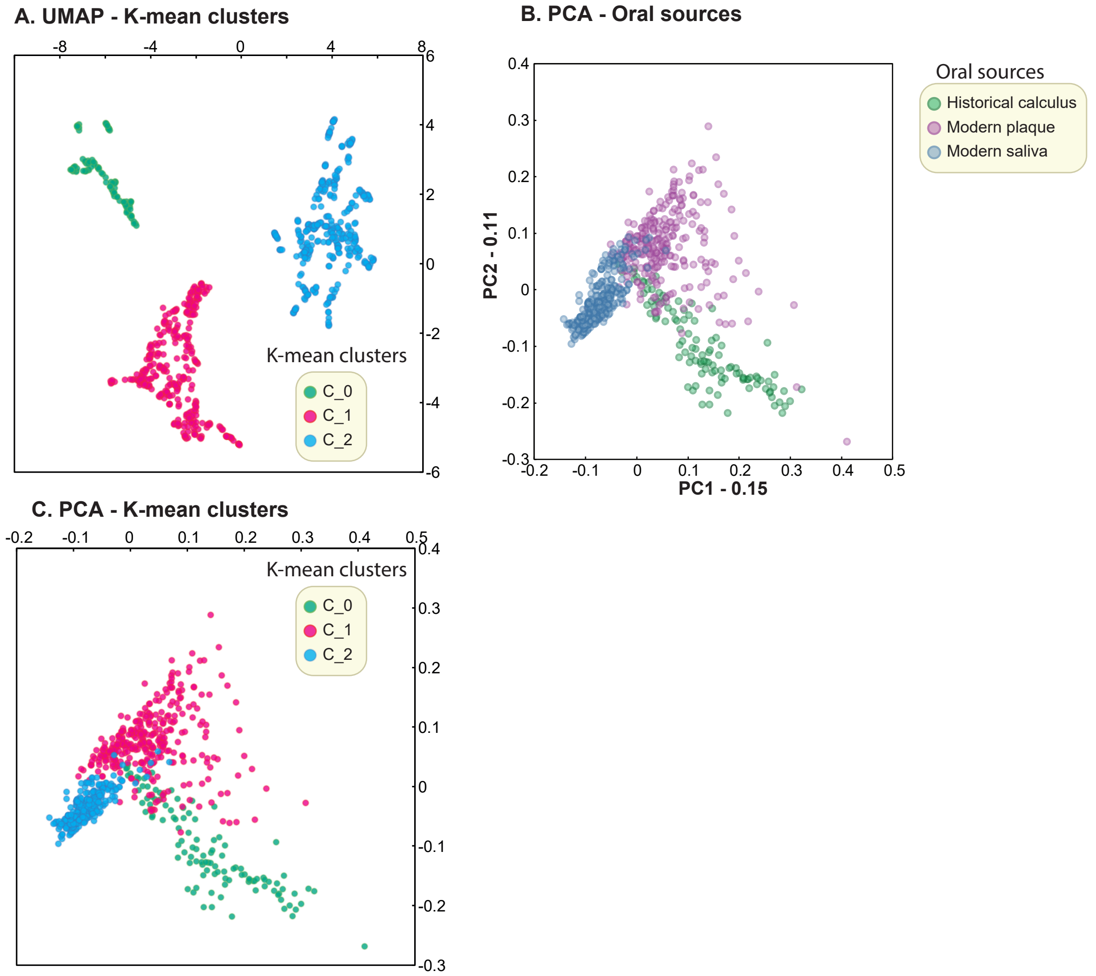


Figure S1. The K-mean clusters of the first three components of the UMAP analyses and their visualization in PCA plots. (A) The visualization of the three clusters in the plane of UMAP components. (B) Plot of the first two components of the PCA analysis. (C) The same plot as (B) with nodes color-coded by K-mean clusters in (A).
