## Supplementary material for "Metagenomics of the modern and historical human oral microbiome with phylogenetic studies on *Streptococcus mutans* and *Streptococcus sobrinus*": Figure S2

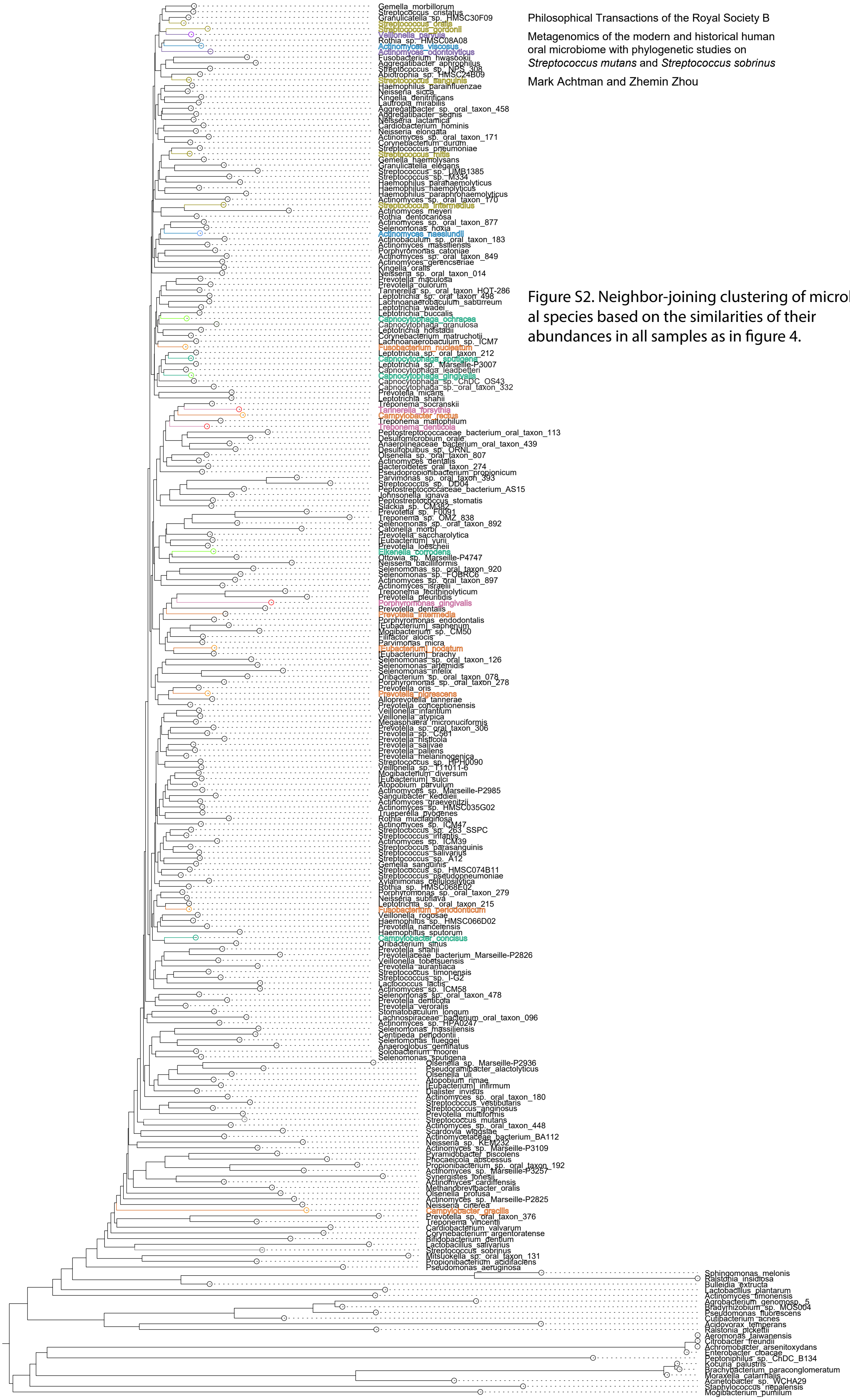


Figure S2. Neighbor-joining clustering of microbial species based on the similarities of their abundances in all samples as in figure 4.
