## Supplementary material for "Metagenomics of the modern and historical human oral microbiome with phylogenetic studies on *Streptococcus mutans* and *Streptococcus sobrinus*": Figure S3

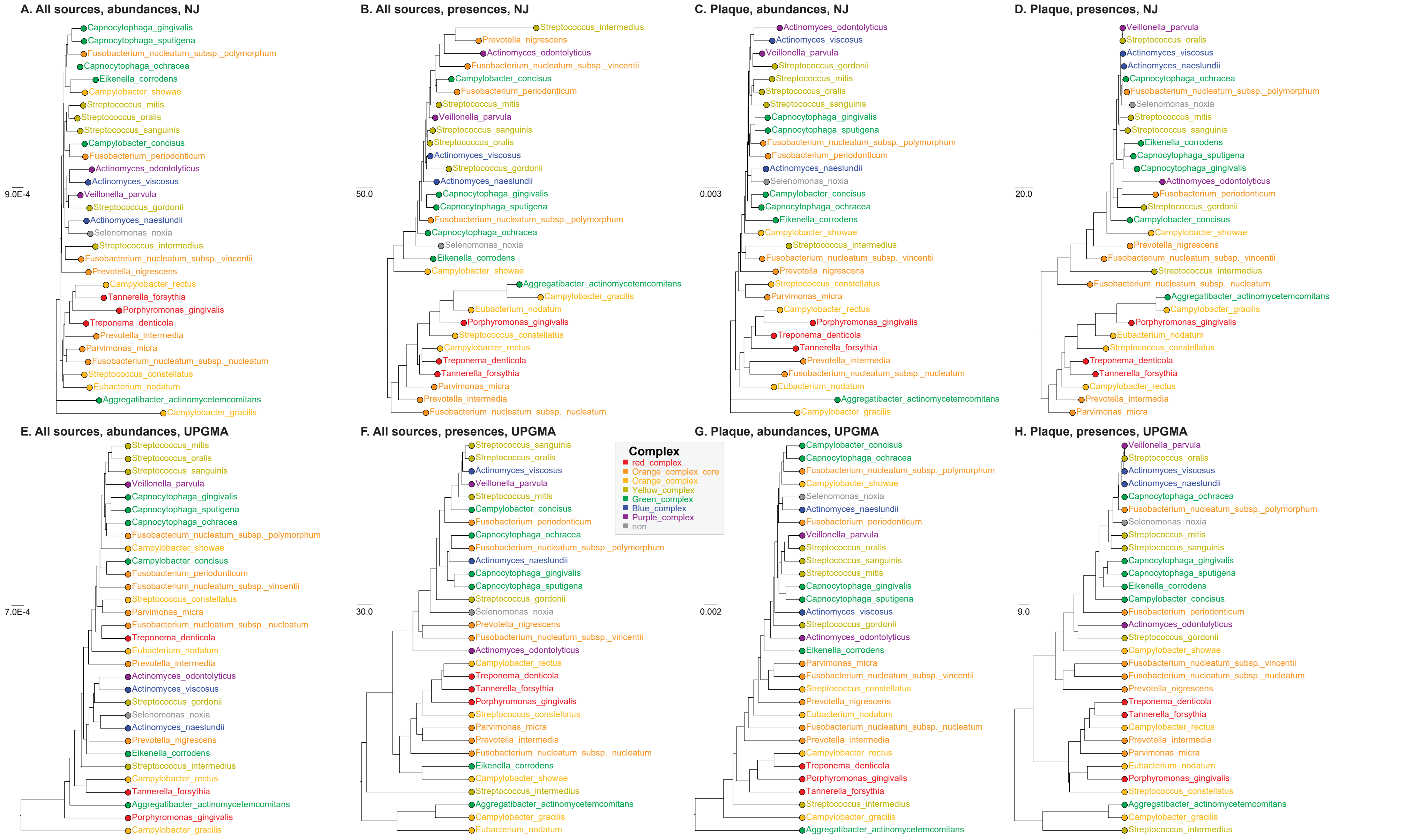


Figure S3. Neighbor-joining and UPGMA clustering of the 28 species described in Socransky et al. based on their abundances or presences in the oral samples.
