## Supplementary material for "Metagenomics of the modern and historical human oral microbiome with phylogenetic studies on *Streptococcus mutans* and *Streptococcus sobrinus*": Figure S4

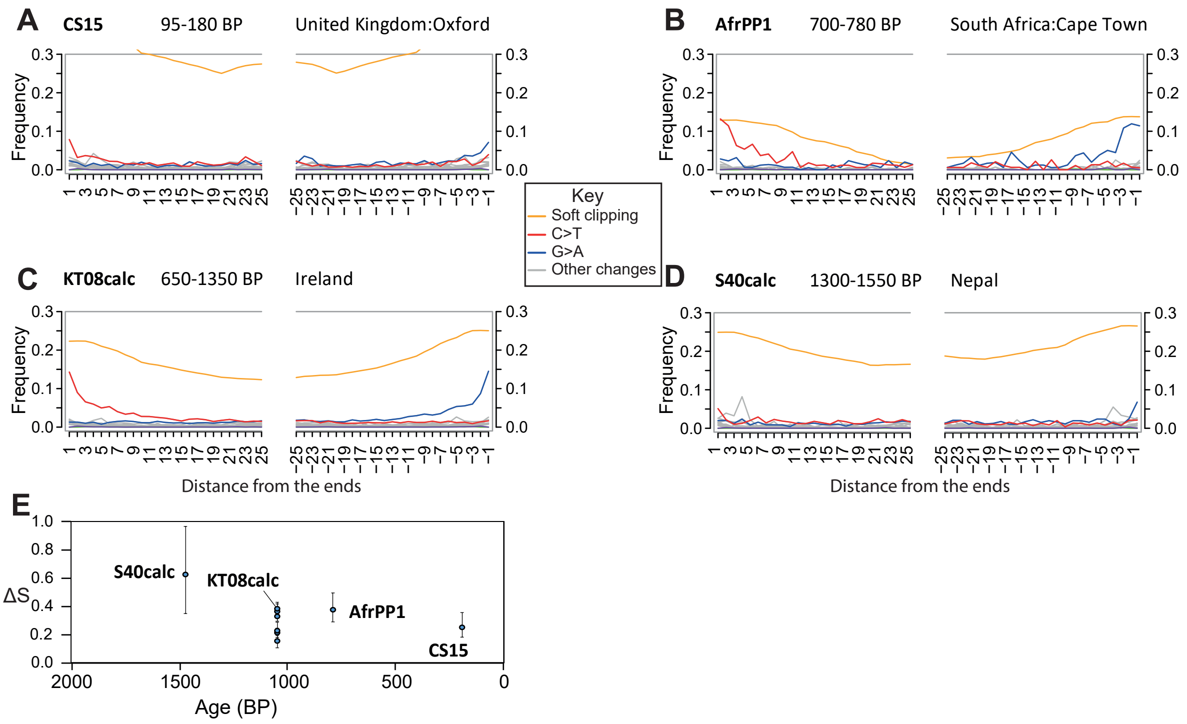


Figure S4. DNA damage at the 5′-end of *S. mutans* sequencing reads from ancient metagenomes. (A-D) MapDamage2 [49] analyses on BAM alignments calculated by Minimap2 [37] of species-specific reads from four selected historical samples against the *S. mutans* UA159 reference genome. Numbers of specific reads were A: 6224; B: 910; C: 26,333; D: 7379. (E) Average MapDamage2 estimates ($\Delta_S$) plus 95% confidence intervals for all ancient metagenomes in which at least 10 *S. mutans* reads were present and at a frequency of ≥0.0001% of all metagenomic reads.
