## Supplementary material for "Metagenomics of the modern and historical human oral microbiome with phylogenetic studies on *Streptococcus mutans* and *Streptococcus sobrinus*": Figure S5

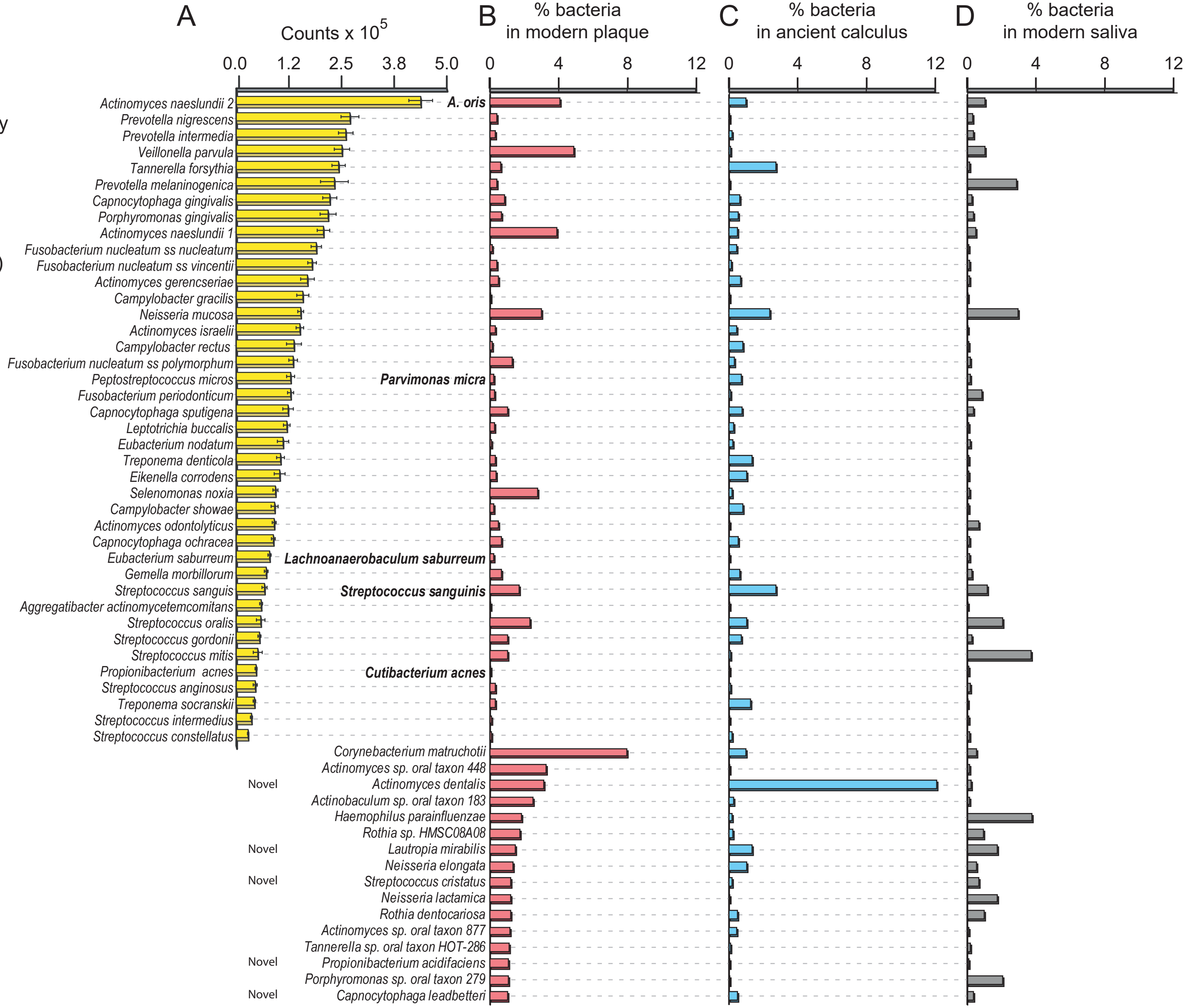


**Figure S5.** A comparison of species abundances between classical checker-board DNA-DNA hybridization and SPARSE. (A) Mean estimated cell counts by DNA-DNA hybridization of 40 species in 20,247 samples from 776 individuals (periodontal disease: 592; healthy: 184). Modified from figure 7 in Socransky and Haffajee, 2005 [28] to include subsequent taxonomic name changes at the right. (B-D) Average percentage of bacterial species in metagenomic samples according to SPARSE in modern dental plaque (B, 307 metagenomes), ancient dental calculus (C, 110) and modern saliva (D, 387). The 40 species at the top correspond to those in part A. Sixteen other species accounted for at least 1% of bacterial reads, and are shown at the bottom. Five of those species were first defined after 1998, and are indicated by 'Novel' at the left.
