## Supplementary material for "Metagenomics of the modern and historical human oral microbiome with phylogenetic studies on *Streptococcus mutans* and *Streptococcus sobrinus*": Figure S6

**Figure S6.** A detailed comparison of species abundances between classical checkerboard DNA-DNA hybridization and SPARSE showing distributions of taxa by periodontal health. A). Data from electronic supplementary material, figure S5 except that the SPARSE data were supplemented with 10 additional metagenomes from dental calculus from patients with dental caries in Spain (6 with periodontal disease and 4 who were healthy) [24]. The sources of the metagenomes are subdivided by health of the donors. B) Breakdown of numbers of metagenomes by source. Health assignments for ancient calculus are according to Velsko *et al.*, 2019.

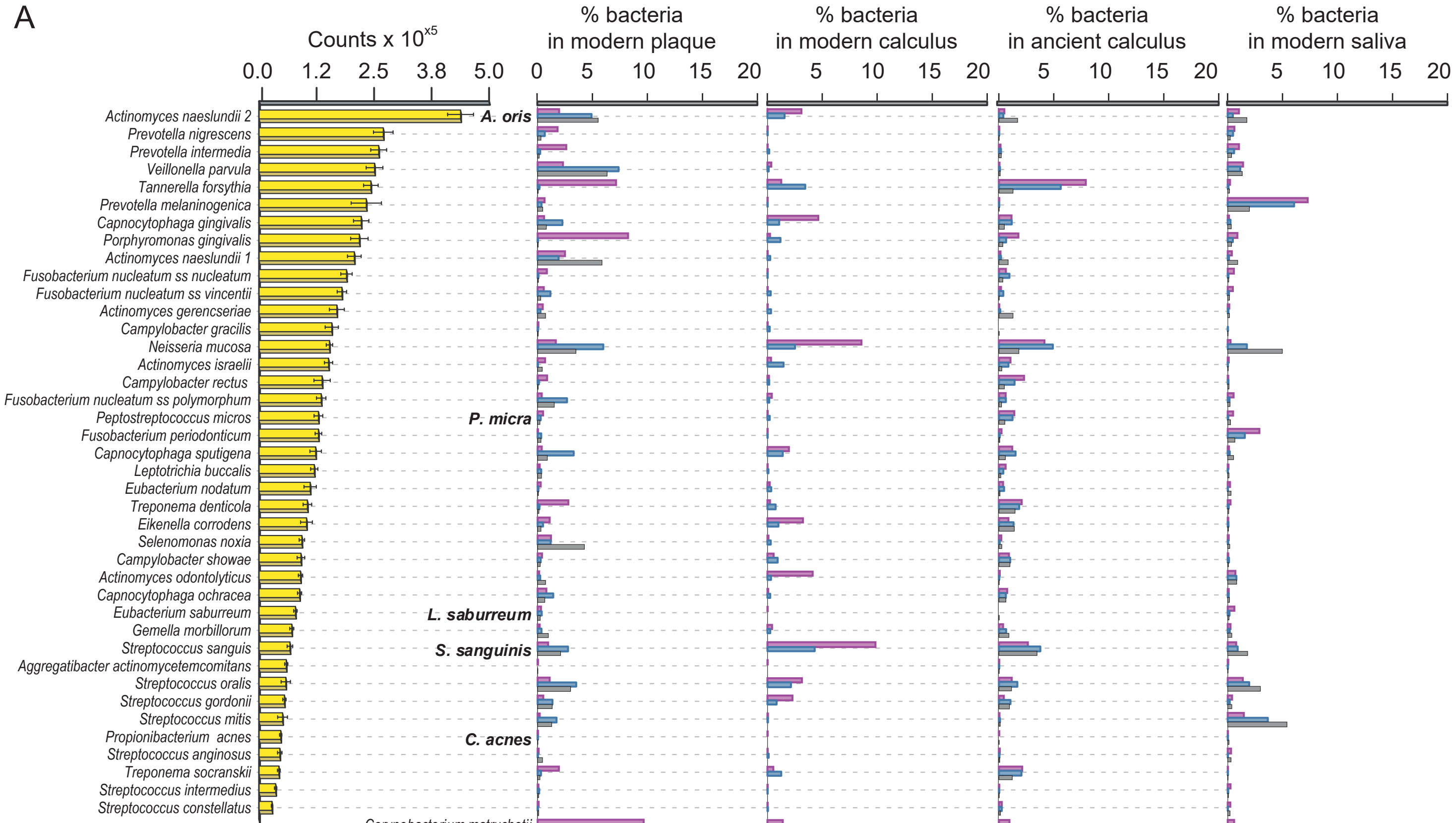


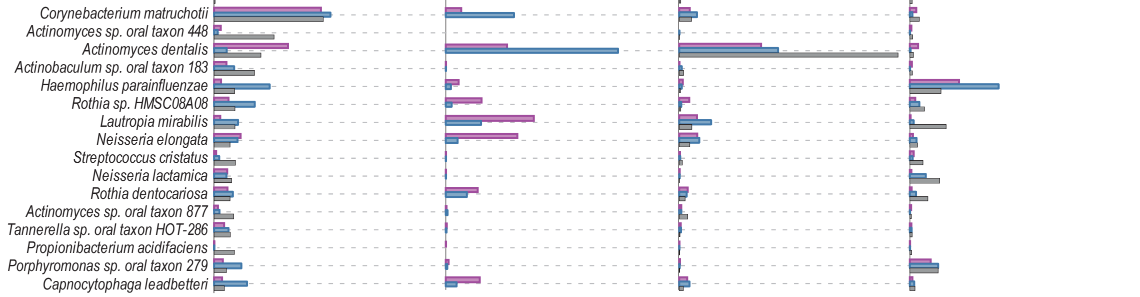


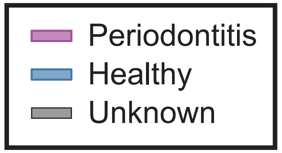


B

| Sample sources | No. samples |
|---|---|
| **Modern plaque:** | |
| Periodontitis | 29 |
| Healthy | 53 |
| Unknown | 225 |
| **Modern calculus:** | |
| Periodontitis | 6 |
| Healthy | 4 |
| **Ancient calculus:** | |
| Periodontitis | 18 |
| Healthy | 26 |
| Unknown | 66 |
| **Modern saliva:** | |
| Periodontitis | 10 |
| Healthy | 139 |
| Others | 238 |
