## Supplementary material for "Metagenomics of the modern and historical human oral microbiome with phylogenetic studies on *Streptococcus mutans* and *Streptococcus sobrinus*": Figure S7

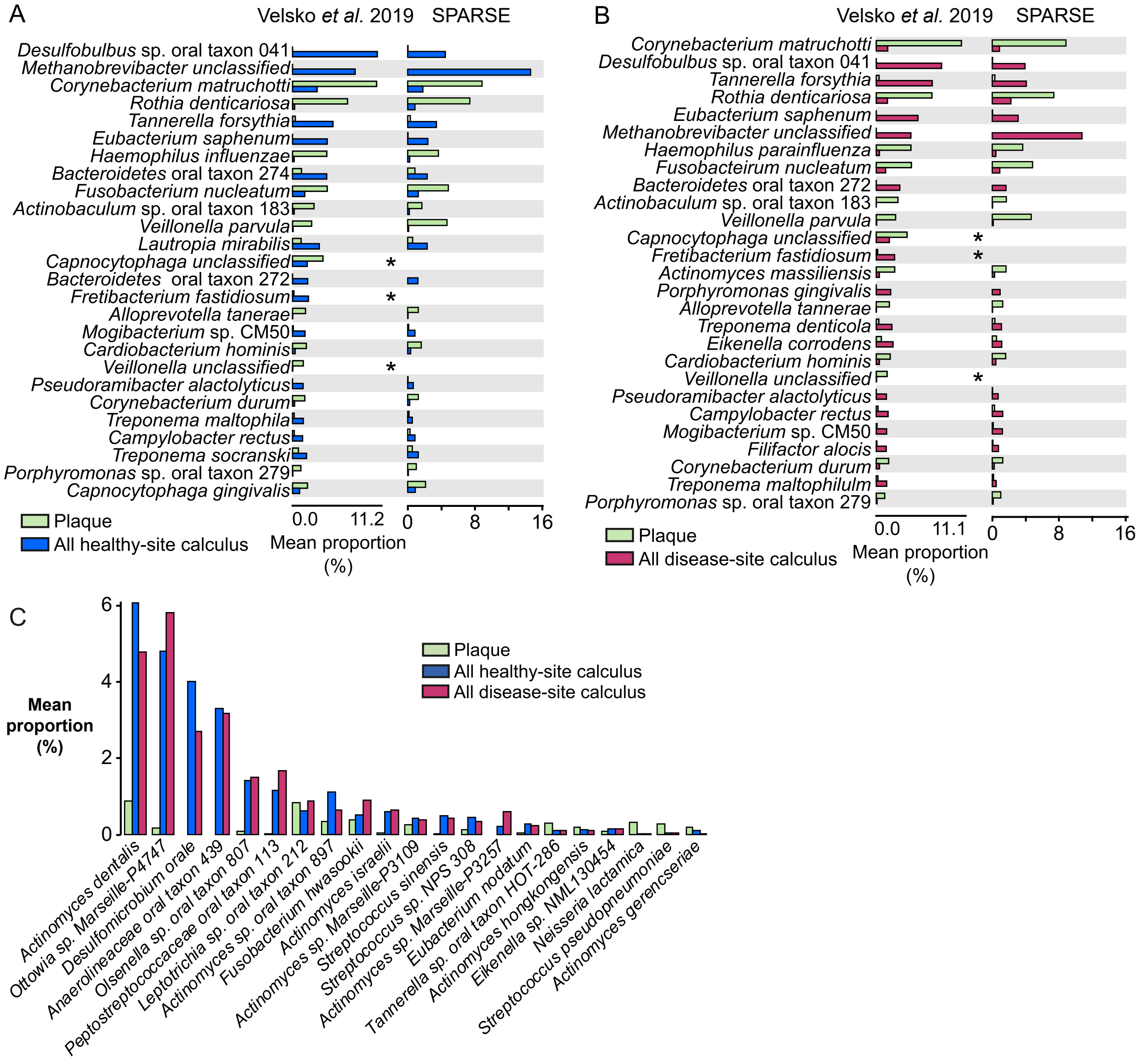


Figure S7. A comparison of species abundances between taxon frequencies within metagenomes in Velsko *et al*., 2019 (which used MetaPhlan2.5.0) and SPARSE. SPARSE results were inversely weighted for the lengths of representative genomes from each of the taxa to allow comparisons with Metaphlan2, which attempts to approximate cell numbers. (A, B) Mean proportions of all microbial cells for abundant species in dental plaque and/or dental calculus. Samples were from donors without periodontal disease (A) or with periodontal disease (B). *Three taxonomic groups are absent from the SPARSE results because the reference genome is of low quality (*F. fastidiosum*) or the Metaphlan2 designations of taxa are above the species level (unclassified *Capnocytophaga, Veillonella*). (C) 21 additional species that were predicted to represent ≥0.1% of cells by SPARSE but not in Velsko *et al.* 2019 although the first five species surpassed the cutoff levels for inclusion by Velsko *et al.* Sources of metagenomes: 20 metagenomes from modern dental plaque: HMP accession codes in Supplemental Table 1, Velsko *et al*., 2019; 10 metagenomes from modern dental calculus: accession code PRJEB31185; 43 metagenomes from ancient dental calculus: accession code PRJEB30331. In parts A and B, the species and results are from Velsko *et al*., 2019, figures 2B and 2E, respectively, with minor modifications.
