## Supplementary material for "Metagenomics of the modern and historical human oral microbiome with phylogenetic studies on *Streptococcus mutans* and *Streptococcus sobrinus*": Figure S8

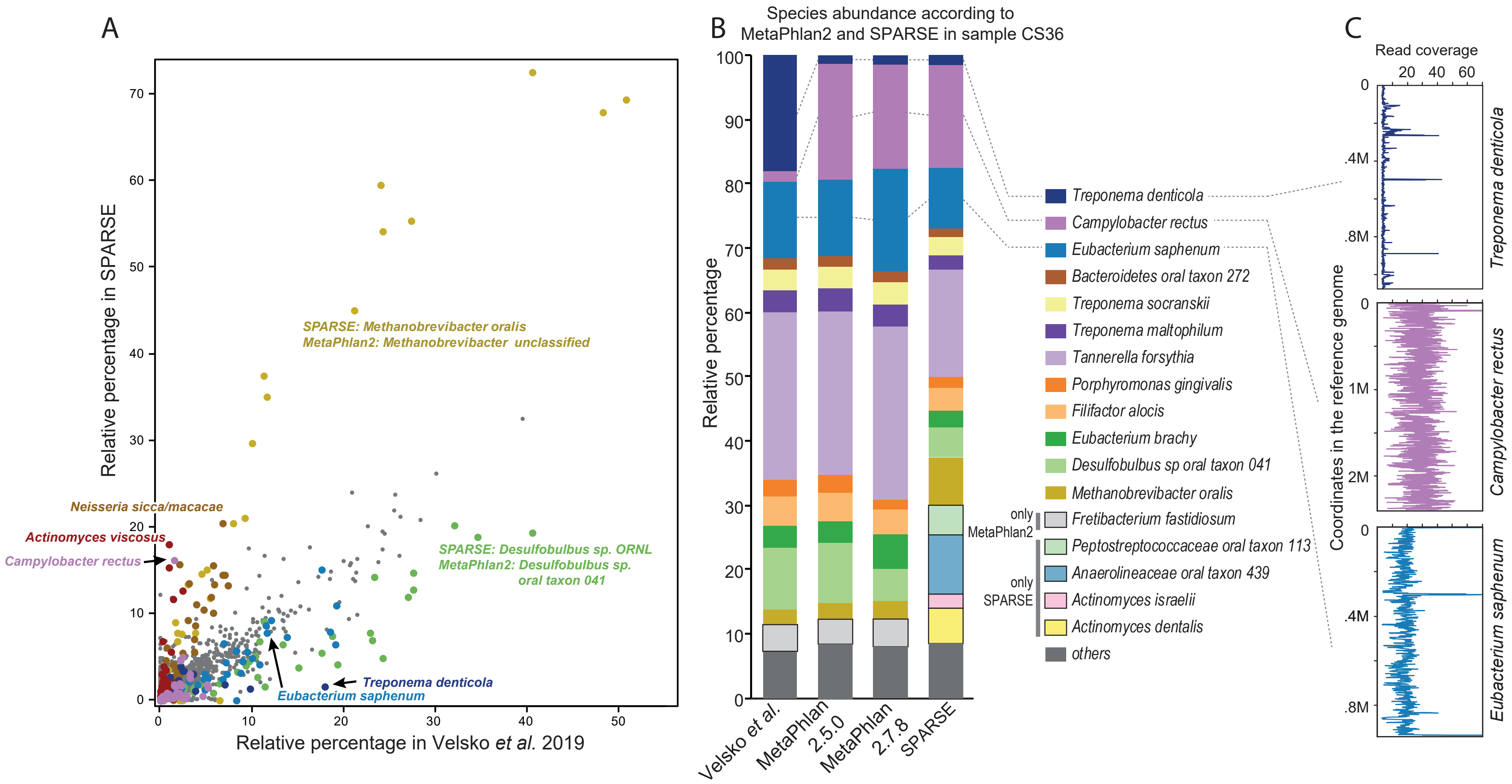


**Figure S8.** A comparison of species abundances between SPARSE, Velsko *et al.* [24] and Metaphlan2. The same species assignments were used as in electronic supplementary material, figure S7, except that reads for CS36 were also assigned to taxa with MetaPhlan V2.5.0 and V2.7.8. (A) Dot plot of the relative percentage of 164 bacterial species that overlapped between supplementary material of Velsko *et al.* 2019 and SPARSE. Two of the species within the genera *Methanobrevibacter* & *Desulfobulbus* are highlighted because they were considered equivalent despite different species designations. Five other species are also highlighted which were off the diagonal, and therefore differentially detected by the two methods. Three arrows indicate data from these species in ancient calculus sample CS36, which was investigated in greater detail in B and C. (B) Abundance of common species in sample CS36 according to Velsko *et al.* 2019 (which used Metaphlan2 V2.5.0), MetaPhlan2 V2.5.0, MetaPhlan2 V2.7.8 (latest version) or SPARSE (after inverse weighting for genome length). Taxa that are specific to SPARSE or Metaphlan2 are bounded by a dark line. (C) Coverage of the reference genomes for three species by metagenomic reads from CS36. Reference genomes: *T. denticola*: GCF_000008185; *C. rectus*: GCF_000174175; *E. saphenum*: GCF_000161975). Reads were mapped to genomes with Minimap2 [37], and the data are the mean coverage for 1 kb windows.
