## Supplementary material for "Metagenomics of the modern and historical human oral microbiome with phylogenetic studies on *Streptococcus mutans* and *Streptococcus sobrinus*": Figure S9

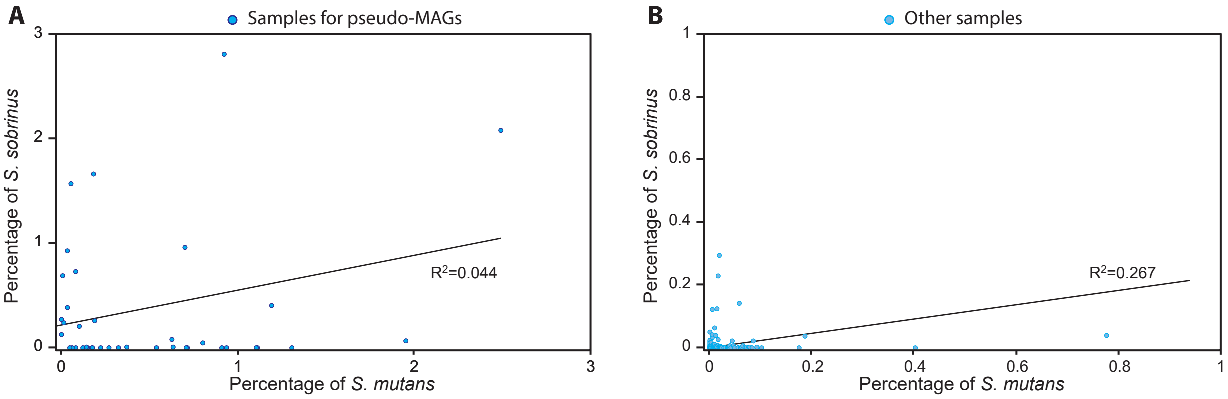


**Figure S9.** Abundances of *S. mutans* (X axis) and *S. sobrinus* (Y axis) in the modern metagenomes that were used to reconstruct pseudo-MAGs (A) and from the low numbers of reads in all other samples (B). The linear regressions (black lines) indicate low correlations between the co-abundances of the two species.
